# Global stability and parameter analysis reinforce therapeutic targets of PD-L1-PD-1 and MDSCs for glioblastoma

**DOI:** 10.1101/2023.05.15.540846

**Authors:** Hannah G. Anderson, Gregory P. Takacs, Duane C. Harris, Yang Kuang, Jeffrey K. Harrison, Tracy L. Stepien

**Affiliations:** Department of Mathematics, University of Florida, Gainesville, FL, USA; School of Mathematical and Statistical Sciences, Arizona State University, Tempe, AZ, USA; Department of Pharmacology and Therapeutics, University of Florida, Gainesville, FL, USA

**Keywords:** immune checkpoint, immunosuppression, mathematical oncology, approximate Bayesian computation, sensitivity analysis

## Abstract

Glioblastoma (GBM) is an aggressive primary brain cancer that currently has minimally effective treatments. Like other cancers, immunosuppression by the PD-L1-PD-1 immune checkpoint complex is a prominent axis by which glioma cells evade the immune system. Myeloid-derived suppressor cells (MDSCs), which are recruited to the glioma microenviroment, also contribute to the immunosuppressed GBM microenvironment by suppressing T cell functions. In this paper, we propose a GBM-specific tumor-immune ordinary differential equations model of glioma cells, T cells, and MDSCs to provide theoretical insights into the interactions between these cells. Equilibrium and stability analysis indicates that there are unique tumorous and tumor-free equilibria which are locally stable under certain conditions. Further, the tumor-free equilibrium is globally stable when T cell activation and the tumor kill rate by T cells overcome tumor growth, T cell inhibition by PD-L1-PD-1 and MDSCs, and the T cell death rate. Bifurcation analysis suggests that a treatment plan that includes surgical resection and therapeutics targeting immune suppression caused by the PD-L1-PD1 complex and MDSCs results in the system tending to the tumor-free equilibrium. Using a set of preclinical experimental data, we implement the Approximate Bayesian Computation (ABC) rejection method to construct probability density distributions that estimate model parameters. These distributions inform an appropriate search curve for global sensitivity analysis using the extended Fourier Amplitude Sensitivity Test (eFAST). Sensitivity results combined with the ABC method suggest that parameter interaction is occurring between the drivers of tumor burden, which are the tumor growth rate and carrying capacity as well as the tumor kill rate by T cells, and the two modeled forms of immunosuppression, PD-L1-PD-1 immune checkpoint and MDSC suppression of T cells. Thus, treatment with an immune checkpoint inhibitor in combination with a therapeutic targeting the inhibitory mechanisms of MDSCs should be explored.

## 1 Introduction

Glioblastoma (GBM) is the most common and aggressive type of primary brain cancer, claiming tens of thousands of lives each year. Within 5 years of diagnosis, less than 10% of GBM patients survive, and most succumb to the tumor within 15 months of diagnosis—even with treatment (Ostrum et al (2015); Fernandes et al (2017)). The current standard of care includes surgical resection followed by radiotherapy and chemotherapy with temozolomide (TMZ) (Stupp et al (2005); Fernandes et al (2017)), but the minimally effective results demonstrate a need to look for new treatment strategies. The highly complex and immune suppressed tumor microenvironment of GBM makes treatment difficult, thus immunotherapy, which has shown to be effective in some cancer types, holds promise (Brown et al (2018)).

In the last decade, an increasing number of immunotherapies have been developed for GBM (Yu and Quail (2021); Bausart et al (2022); Bryukhovetskiy (2022)). Anti-PD-1 is a standard immunotherapy that has proven to be beneficial in treating other cancers, such as melanoma (Postow et al (2015)), Hodgkin’s lymphoma (Ansell et al (2015)), colon (Duraiswamy et al (2013)), cervical (Chung et al (2019)), and non-small cell lung cancer (NSCLC) (Brahmer et al (2015)), although it has failed as a monotherapy in GBM phase III clinical trials (Preusser et al (2015); Reardon et al (2017); Lim et al (2018)). Cancer can evade the immune system by expressing the marker PD-L1, which downregulates the cytotoxic response of activated T cells by binding to their PD-1 receptor. Treatment with anti-PD-1 unmasks the tumor by binding to PD-1, thereby facilitating tumor recognition by T cells and enhancing the immune response. The failure of anti-PD-1 monotherapy in GBM might potentially be explained by additional mechanisms of immune suppression, such as the infiltration of myeloid-derived suppressor cells (MDSCs), which are cells that suppress T cell activity.

Gliomas mediate recruitment of CCR2^+^ MDSCs by releasing chemokines such as CCL2 and CCL7, which are cognate ligands of the CCR2 receptor (Takacs et al (2021, 2022)). Higher expression of CCL2 and CCL7, as well as CCR2, within the tumor microenvironment all correlate with a worse prognosis for gliomas (Korbecki et al (2020); Chang et al (2016)). Once the MDSCs enter the tumor microenvironment, they suppress T cells through a variety of mechanisms including, but not limited to, inducing apoptosis in activated T cells (Saio et al (2001)), producing enzymes (such as arginase) which metabolize amino acids needed for T cell proliferation (Condamine and Gabrilovich (2011)), and secreting other factors which modulate the immune response (NO, TGF-*β*, TNF-*α*, H_2_O_2_) (Monu and Frey (2012); Markowitz et al (2017)). These mechanisms, along with MDSC promotion of angiogenesis (Vetsika et al (2019)), lead to rapid tumor progression. Treating with a CCR2 antagonist aids the immune response by reducing MDSC recruitment and thus sequestering the MDSC population to the bone marrow. Flores-Toro et al (2020) reported improved survival of two murine glioma models (KR158 and 005 GSC) with combination treatment of a CCR2 antagonist (CCX872) and anti-PD-1.

Although MDSCs play a significant role in glioma progression, there are few published mathematical models of tumor–immune dynamics that incorporate these cells. Allahverdy et al (2019) presents a discrete agent-based model of MDSCs in the context of a general tumor and simulates treatment with the chemotherapeutic agent 5-fluorouracil (5-FU). Similar to Allahverdy et al (2019), the ODE model of Shariatpanahi et al (2018) for a general tumor also simulates treatment with 5-FU. Further, they include treatment with L-arginine since the model considers production of arginase I as the primary mechanism by which MDSCs suppress T cells. Liao et al (2014) highlights a mechanism of MDSC recruitment by developing a PDE model which focuses on the role of interleukin-35 (IL-35) in MDSC recruitment and tumor growth. Lai et al (2018) used a PDE model of breast cancer to determine the influence of the immune checkpoint inhibitor, anti-CTLA-4, on M2 macrophages within the tumor site. Although they did not specifically model MDSCs, their work implied that these conclusions apply to MDSCs because a subset of monocytic MDSCs differentiate into M2 macrophages. Kreger et al (2023) developed a stochastic delay differential equations model of a metastasizing breast cancer to evaluate MDSC influence on tumor progression. Although the model is fitted to patient response data to immune checkpoint inhibitors, the model itself does not include a mechanism to represent an immune checkpoint. Each of these models include MDSCs and provide a foundation to incorporate the PD-L1-PD-1 immune checkpoint. Furthermore, we aim to develop a model specifically for GBM instead of a general tumor.

Mathematical models of tumor–immune dynamics range from modeling general tumors (Lai and Friedman (2017); Nikolopoulou et al (2018); Eftimie et al (2011); Mahlbacher et al (2019); Shi et al (2021); Radunskaya et al (2018); Khyat and Jang (2022)) to specific cancers including GBM, non-small lung cancer (NSCLC), melanoma, and breast cancer (Storey et al (2020); Yu and Jang (2019); Butner et al (2021); Lai et al (2018); Banerjee et al (2015); Özköse et al (2022); Mirzaei et al (2021); Khajanchi and Banerjee (2017); Khajanchi (2021); Jafarnejad et al (2019); Perlstein et al (2019); Bitsouni and Tsilidis (2022)). Similar to Lai et al (2018), Yu and Jang (2019) modelled the CTLA-4 immune checkpoint and included anti-CTLA-4 therapy, but restricted their focus to a four equation ODE model of tumor cells, CD4^+^ T cells, IFN-*γ*, and CTLA-4. Two years earlier, Lai and Friedman (2017) modelled a different immune checkpoint, the PD-L1-PD-1 complex, by developing a PDE model of a general tumor which included additional cells and molecules like dendritic cells, CD4^+^/CD8^+^ T cells, IL-2, and IL-12. Their model was used to simulate treatment with anti-PD-1 and a cancer vaccine, GVAX. This work was later reduced to an ODE model by Nikolopoulou et al (2018) which focused on tumor and T cells with intermittent and continuous anti-PD-1 treatment. Shi et al (2021) and Nikolopoulou et al (2021) extended this work by establishing the global dynamics of the model along with a generalized version. Storey et al (2020) applied these models to GBM by estimating parameters from in vivo murine experimental data. Their ODE model divided the immune compartment into the categories of innate and adaptive immune cells and modeled treatment with anti-PD-1 and an oncolytic viral therapy. Radunskaya et al (2018) deviated from modeling the typical immune checkpoint inhibitor, anti-PD-1, by considering treatment with anti-PD-L1 in their model of general tumor-immune dynamics, which compartmentalized interactions to be within the spleen, blood, and tumor site. While each of these previous papers focused on a specific immune checkpoint, Butner et al (2021) developed an ODE model which was not necessarily specific to the type of immune checkpoint or the type of tumor. Using this model along with patient data, Butner et al (2021) predicted the long-term tumor burden for a variety of patients being treated with different immune checkpoint inhibitors.

In this paper, we develop a GBM-specific ODE model of cancer and T cell interactions by incorporating the PD-L1-PD-1 immune checkpoint along with tumor recruitment of MDSCs. Parameter values are estimated from the literature and via comparison with experimental in vivo murine data.

Our paper unfolds as follows: in Section 2, we describe the GBM-immune dynamics model. In Section 3, we perform equilibrium and solution analysis and emphasize these findings through bifurcation analysis. Section 4 analyzes the parameter space by implementing the Approximate Bayesian Computation (ABC) method used in conjunction with the extended Fourier Amplitude Sensitivity Test (eFAST). Numerical simulations alongside experimental data are also provided. In Section 5, we conclude with a discussion of our results and future directions.

## 2 Mathematical Model

The glioblastoma (GBM)–immune model focuses on the dynamics within the glioma microenvironment and is based on the previous work of Nikolopoulou et al (2018), Storey et al (2020), and Shariatpanahi et al (2018). A diagram of the biological interactions implemented is illustrated in Fig. 1, and Table 1 lists the biological meaning of the model’s parameters with their respective units and representative range of values.

**Table 1:**
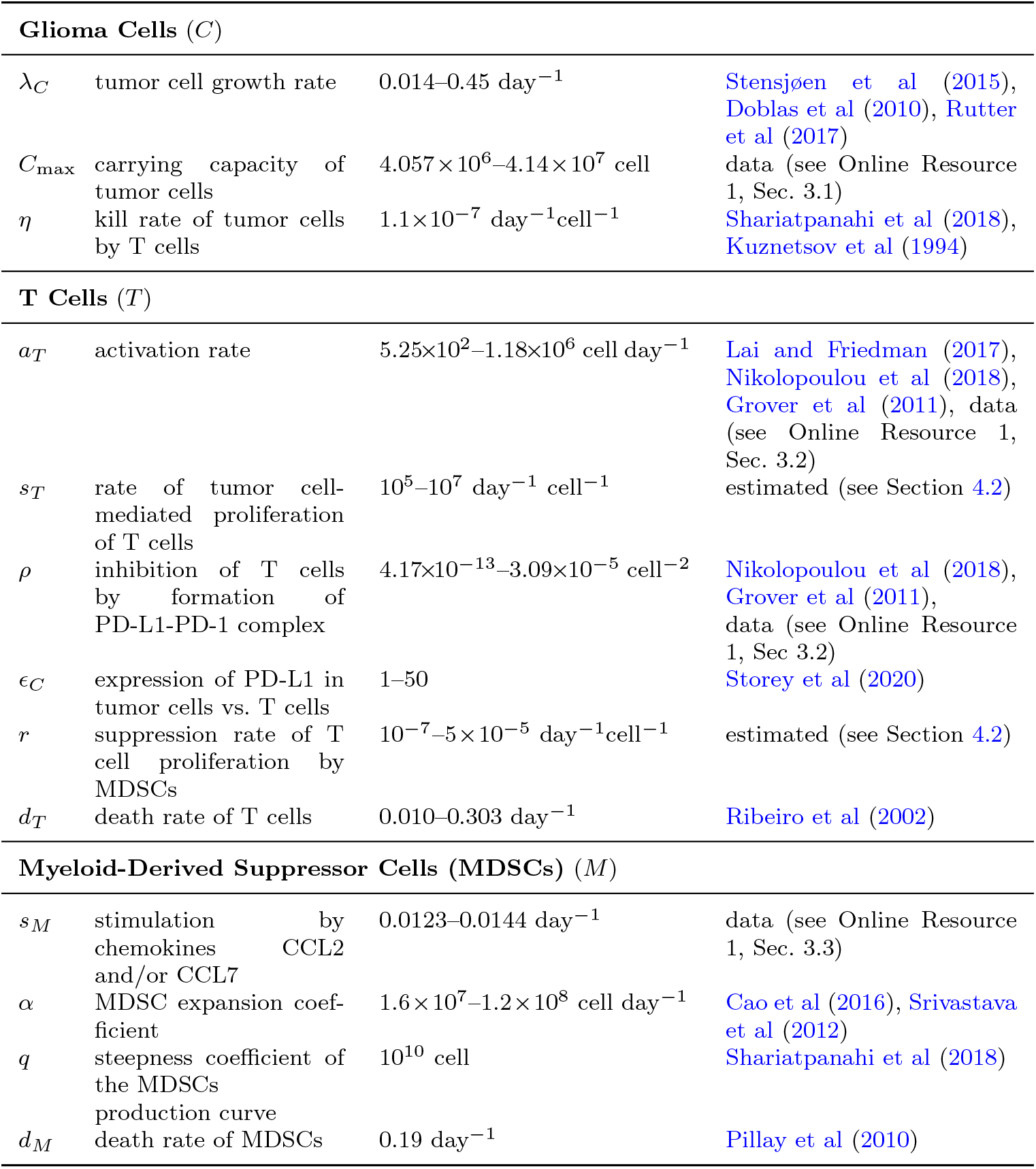
Model parameters of the glioblastoma (GBM)–immune dynamics model (1).

**Fig. 1:**
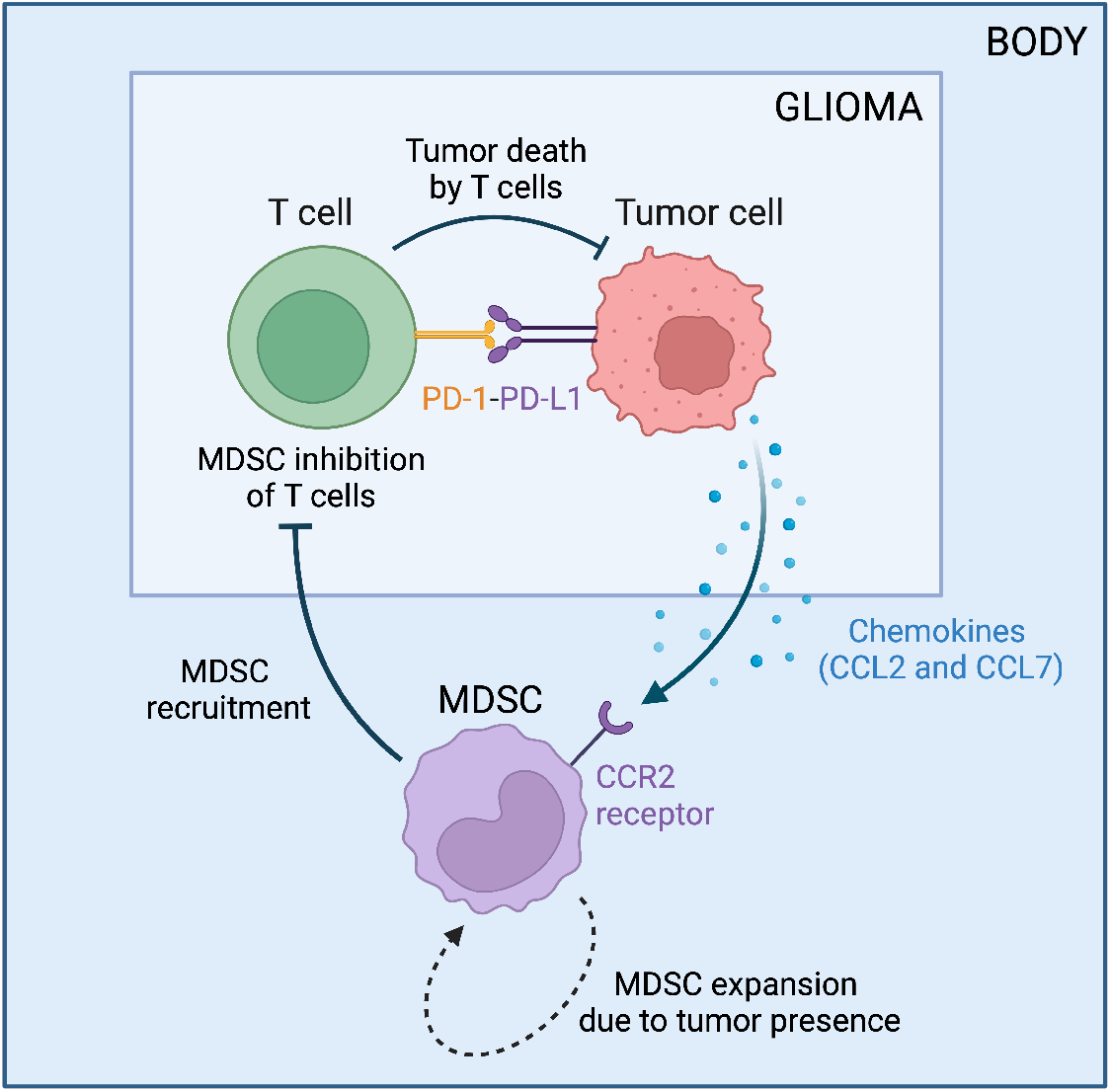
Tumor-immune interactions in glioblastoma. Flowchart created with BioRender.com.

Letting *C* be the number of tumor cells, *T* the number of activated T cells (all cells that have the CD3^+^ marker, which includes both CD4^+^ and CD8^+^ T cell subsets), and *M* be the number of myeloid-derived suppressor cells (MDSCs), we arrive at the following system of equations:

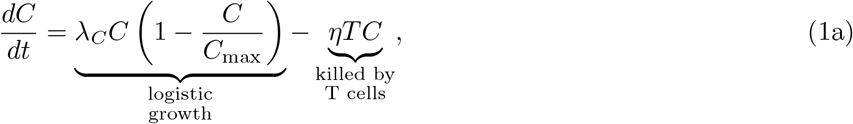

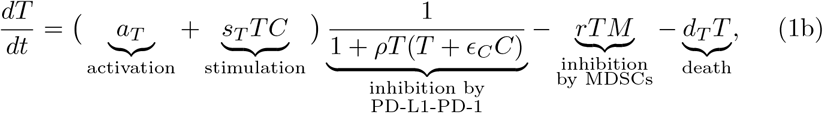

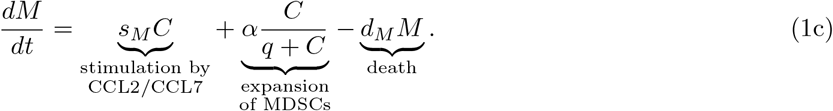

In equation (1a), we assume the tumor cells exhibit logistic growth with a carrying capacity of *C*_max_. T cells, which are the main drivers of tumor cell death, kill glioma cells at a rate of *η*, as in Nikolopoulou et al (2018).

Equation (1b) is of the same form as in Lai and Friedman (2017), Nikolopoulou et al (2018), and Storey et al (2020). The first term represents T cells activation at a constant rate *a*_*T*_, potentially due to the cytokine IL-12 which is explicitly represented in the previous models, and the recruitment (or stimulation) of T cells to the tumor site by the presence of a glioma at a rate *s*_*T*_ . Once in the glioma microenvironment, T cells are exhausted by the formation of the PD-L1-PD-1 complex, which binds PD-L1 present on glioma cells and T cells with PD-1 on T cells. Assuming (*T* + *ϵ*_*C*_*C*) characterizes the level of free PD-L1 in the tumor microenvironment, the PD-L1-PD-1 complex is represented by *T*(*T* + *ϵ*_*C*_*C*) since only T cells express PD-1. As T cells and tumor cells increase, more PD-L1-PD-1 complexes form, thus decreasing the overall fraction to represent T cell inhibition. The parameter *ρ* results from combining multiple parameters in Lai and Friedman (2017), Nikolopoulou et al (2018), and Storey et al (2020) due to their structural non-identifiability. The remaining terms of (1b) represent T cells suppressed/deactivated by MDSCs (Gabrilovich and Nagaraj (2009)) by a rate *r* and T cells dying at a rate *d*_*T*_ .

In equation (1c), the first term represents glioma recruitment of MDSCs to the microenvironment by secreting the chemokines CCL2 and CCL7, which are ligands for the CCR2 receptor expressed by MDSCs. This results in a chemotactic effect, drawing MDSCs from the bone marrow to the glioma microenvironment, where here we are specifically focusing on monocytic M-MDSCs as opposed to granulocytic PMN-MDSCs. Since these chemokines are produced by glioma cells, this recruitment increases as glioma cells increase. We assume that the presence of a tumor also results in the expansion of splenic MDSCs (Liu et al, 2007, Fig. 2), using the same fractional term as the model of Shariatpanahi et al (2018) in the second term of (1c), thus indirectly causing more MDSCs to accumulate in the tumor. The last term represents MDSCs dying at a rate *d*_*M*_ .

**Fig. 2:**
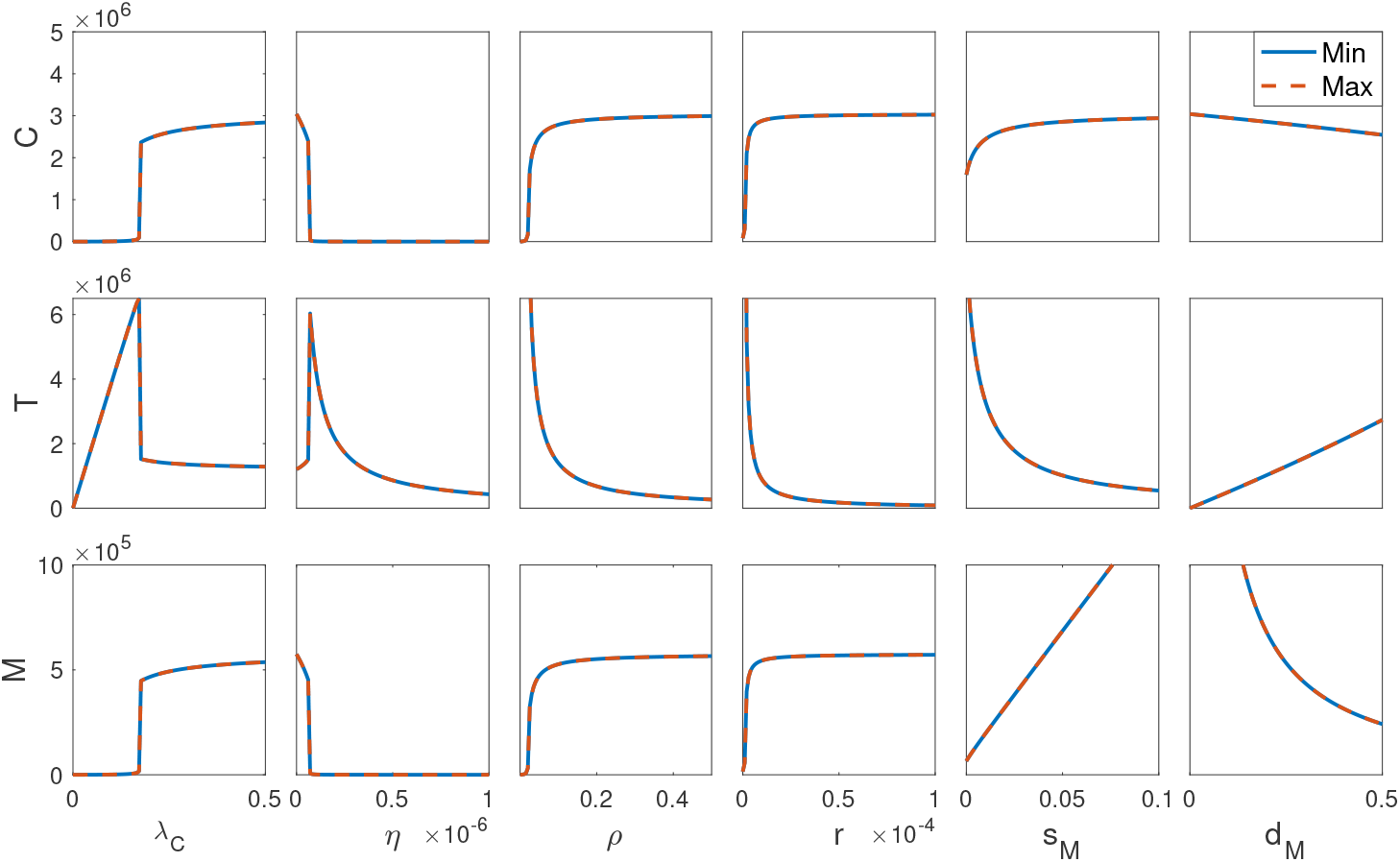
Bifurcation diagrams for the tumor growth rate (*λ*_*C*_), T cell kill rate of tumor cells (*η*), T cell inhibition rate by PD-L1-PD-1 (*ρ*) and MDSCs (*r*), and MDSC recruitment (*s*_*M*_) and death rate (*d*_*M*_). The minimum/maximum tumor (*C*), T cell (*T*), and MDSC (*M*) cell counts are shown in solid blue/dashed red, respectively. While all parameters can decrease the immune response, only varying *λC, η, ρ*, and *r* can eliminate the tumor cells, indicating their potential as therapeutic targets.

Our model is distinct from previous models in that it combines both the PD-L1-PD-1 immune checkpoint, such as incorporated in Lai and Friedman (2017), Nikolopoulou et al (2018), and Storey et al (2020), and MDSCs, such as in Shariatpanahi et al (2018), and it additionally was developed specifically for GBM. Though Storey et al (2020) also focused on GBM, while the other studies were for general tumors or tumors different than GBM, their study included oncolytic viral therapy treatment. Shariatpanahi et al (2018) differs from our model in that MDSC dynamics are modeled in the spleen instead of the tumor site and they incorporated chemotherapy treatment. Here, we integrate an immune checkpoint mechanism and MDSCs to study each of these immunotherapy targets in GBM.

## 3 Equilibrium and Solution Analysis

We describe the equilibria and stability of system (1) in this section to verify that our model is biologically reasonable and to determine suitable target parameters for treatment. Since the functions on the right hand side of the equations in system (1) are continuously differentiable, solutions exist and are unique by standard ordinary differential equations theory.

We begin by confirming that our system will neither veer into negative cell counts nor increase to infinity over time but will be realistically bounded. First, we show that solutions with positive initial conditions remain positive for all time.

### Theorem 1

(Positivity) *Solutions of* (1) *that start positive remain positive*.

*Proof* Assume *C*(*t*_0_) > 0, *T*(*t*_0_) > 0, and *M*(*t*_0_) > 0 for some initial time *t*_0_. Without loss of generality, assume *t*_0_ = 0.

In order to have *T*(*t*) ≤ 0 for some *t* > 0, it is necessary to have *dT/dt* ≤ 0 for *T* = 0. However, when *T* = 0, (1b) becomes *dT/dt* = *a*_*T*_ > 0. Thus, *T*(*t*) > 0 for all *t* > 0.

Assuming that *C* and *T* exist on some interval [0, *t*) for *t* > 0, when we integrate (1a), we arrive at

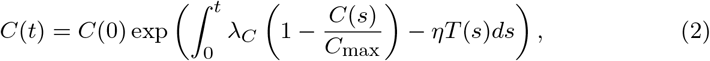

which is always positive since *C*(0) > 0.

Lastly, when *M* = 0, (1c) becomes

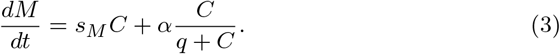

Since we proved that *C*(0) > 0 implies *C*(*t*) > 0 for all *t* > 0 and because all the parameters are positive, we conclude that for *M* = 0, *dM/dt* > 0. Thus, *M*(*t*) > 0 for all *t* > 0. □

Next, we show that solutions of system (1) with positive initial conditions are bounded.

### Theorem 2

(Boundedness) *Solutions of* (1) *that start positive are bounded*.

*Proof* Assume *C*(*t*_0_) > 0, *T*(*t*_0_) > 0, and *M*(*t*_0_) > 0 for some initial time *t*_0_. Without loss of generality, assume *t*_0_ = 0.

Since all parameters are positive and *C* and *T* are positive by Theorem 1, by comparison we can bound (1a) by

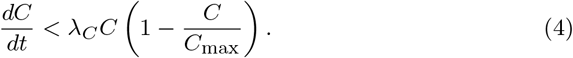

Since logistic growth is bounded, *C*(*t*) is bounded as well. Specifically,

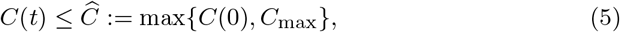

for all *t* ≥ 0.

For the T cells, note that (1b) can be bounded as

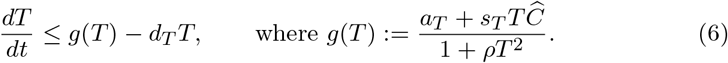

*g*(*T*) achieves an absolute maximum at

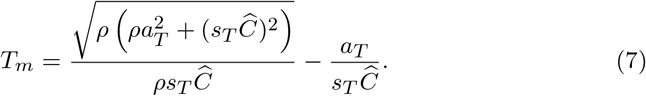

Hence, (6) can be further bounded as

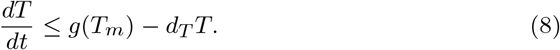

Using a standard comparison argument,

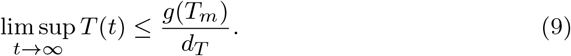

Thus, for all *t* ≥ 0,

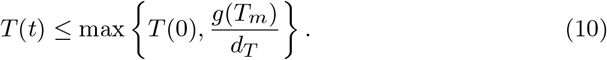

For the MDSCs, since *C* is bounded (5), (1c) can be bounded as

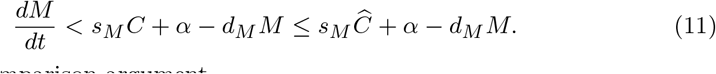

By a standard comparison argument,

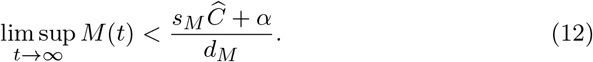

Therefore, for all *t* ≥ 0,

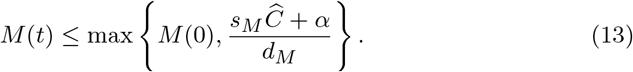

Hence, system (1) is bounded. □

The system (1) admits two categories of fixed points: tumor-free equilibria of the form 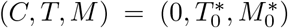 and tumorous equilibria of the form 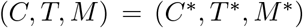. We first show that there exists one unique tumor-free equilibrium.

### Theorem 3

(Uniqueness of tumor-free equilibrium) *The system* (1) *has a unique tumor-free equilibrium* 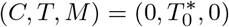, *where* 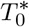 *is increasing with respect to a*_*T*_ *and decreasing with respect to d*_*T*_ *and ρ*.

*Proof* Assume that there is no tumor present, i.e., *C* = 0. Setting *dM/dt* = 0 in (1c) implies that 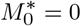.

When *dT/dt* = 0, (1b) implies that 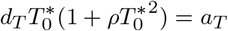 . Let

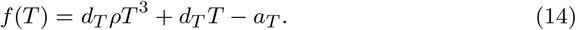

According to Descartes’ rule of signs, the number of sign changes in front of the coefficients of a polynomial corresponds to the number of positive zeros. Since there is exactly one sign change because all the parameters are positive, there is exactly one positive zero, 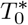. Thus, a biologically relevant tumor-free equilibrium 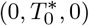 exists and is unique.

Note that as *a*_*T*_ increases, the graph of *f*(*T*) shifts down, resulting in a larger 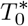. As *d*_*T*_ and *ρ* increase, *f′*(*T*) = 2*d*_*T*_ *ρT* ^2^ + *d*_*T*_ increases for *T* > 0, which results in a narrowed curve and thus a smaller 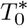.

Lastly, when we solve for 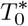 using the cubic formula, we obtain

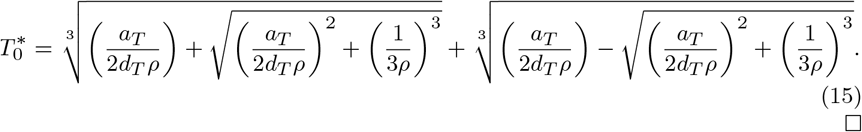

□

Next, we determine conditions for the stability of the tumor-free equilibrium 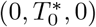.

### Theorem 4

(Stability of tumor-free equilibrium) *When* 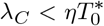, *the tumor-free equilibrium*, 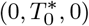, *is locally asymptotically stable. However, when* 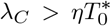 *is a saddle point*.

*Proof* The Jacobian evaluated at the tumor-free equilibrium is

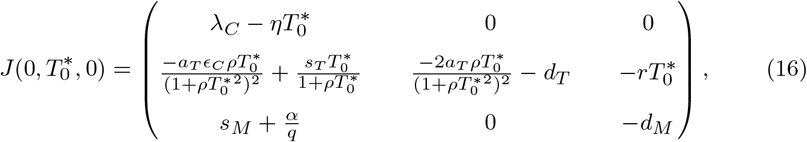

and the eigenvalues are

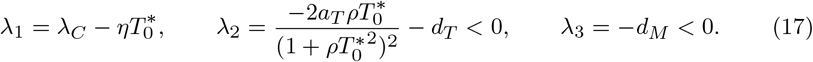

Therefore, when 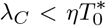, the tumor-free equilibrium 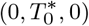 is locally asymptotically stable. However, when 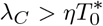, so the tumor-free equilibrium is a saddle point. □

Biologically, the condition 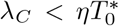 guaranteeing local stability of the tumor-free equilibrium indicates that T cells kill cancer cells faster than the cancer cells can multiply during the onset of tumor initiation. As for treatment, the worst patient scenario would be a saddle tumor-free equilibrium, as this would suggest that the patient will most likely relapse regardless of their proximity to the disease-free condition. Thus, by varying *λ*_*C*_, *η*, or 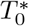 (15) (which depends on *a*_*T*_, *d*_*T*_, and *ρ*), a patient’s tumor-free equilibrium could shift from a saddle to a locally stable point. These parameters would likely be affected in an immune-compromised individual, and in particular, treatment with an immune checkpoint inhibitor to decrease *ρ* could be an effective strategy.

In Appendix A, we establish conditions for the uniqueness (Theorem 6) and local stability (Theorem 7) of the tumorous equilibrium. We conclude our discussion here by determining conditions for the global stability of the tumor-free equilibrium 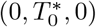.

### Theorem 5

(Global stability of tumor-free equilibrium) *The system* (1) *has a globally stable tumor-free equilibrium when λ*_*C*_ < *ηβ, where* 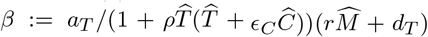 *and* 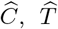 *and* 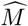*are the upper bounds for glioma cells* (5), *T cells* (10), *and MDSCs* (13), *respectively, as determined in Theorem 2*.

*Proof* From Theorems 1–2, we have that *C*(*t*), *T*(*t*), and *M*(*t*) are bounded above with upper bounds denoted by 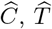 and 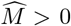, respectively, and below by 0. By these bounds,

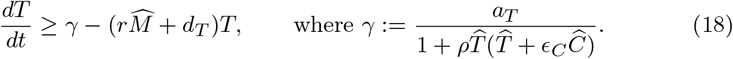

Thus, using a standard comparison argument,

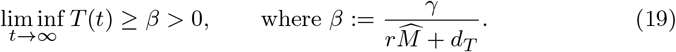

If *λ*_*C*_ < *ηβ*, then for all *s* > 0 sufficiently large,

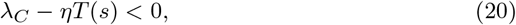

which, taking the limit of (2), implies that

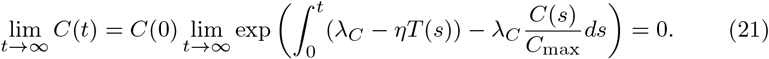

Bounding *T*(*t*) in the tumor cell equation (1a), we have

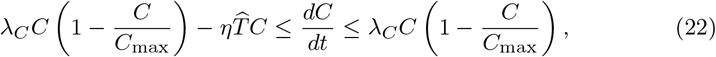

which, taking the limit as *t* → ∞, implies that *dC/dt* → 0. Now, using (21),

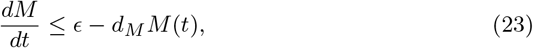

where *ϵ* → 0 as *t* → ∞. A standard comparison argument reveals that

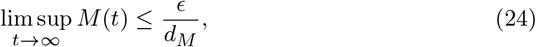

for any *ϵ* > 0. Thus, since *M*(*t*) is nonnegative for all *t* ≥ 0, *M*(*t*) → 0 as *t* → ∞, and hence, *dM/dt* → 0 as *t* → ∞.

Since *T* is bounded above and below and since *C*(*t*) → 0 and *M*(*t*) → 0 as *t* → ∞, the limiting equation of (1b) is *dT/dt* = *G*(*T*), where

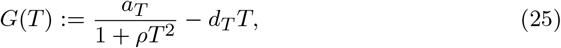

and we note that 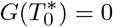. Since

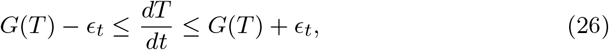

where *ϵ*_*t*_ → 0 as *t* → ∞, using a comparison argument,

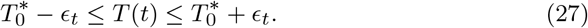

Thus, 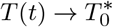as *t* → ∞, and 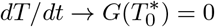 as *t* → ∞.

This allows us to conclude that the tumor-free equilibrium point, 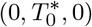, is globally stable in 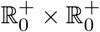 under the condition that *λ*_*C*_ < *ηβ*. □

The best patient scenario is one in which the tumor will tend toward the tumor-free equilibrium regardless of size. While satisfying the global stability condition in Theorem 5 is unlikely in reality, as we shall see in Section 4.3, the condition provides additional targets for treatment, such as *ϵ*_*C*_, *r*, and *C*_max_. These parameters could be targeted through treatment with anti-PD-1/PD-L1, L-arginine, and surgical resection, respectively. Through parameter analysis in Section 4, we will narrow our search for therapeutic targets.

### 3.1 Bifurcation Analysis

We validate the tumor-free equilibrium global stability conditions (Theorem 5) with a bifurcation analysis. Fig. 2 indicates that the tumor growth rate (*λ*_*C*_), T cell kill rate of tumor cells (*η*), and T cell inhibition rate by PD-L1-PD-1 (*ρ*) and MDSCs (*r*) are reasonable targets for treatment, since shifting these parameters can result in the system tending toward the tumor-free equilibrium. Varying the remaining parameters cannot considerably relieve the tumor burden (Fig. S1 (Online Resource 1, Sec. 5)) and thus are unlikely to be advantageous treatment targets.

Regarding MDSC infiltration and suppression, Fig. 2 suggests that targeting MDSC recruitment, death, and their inhibition of T cells can increase the cytotoxic immune response; however, only by directly targeting immune suppression by MDSCs, *r*, can tumor cells be reduced to zero. Thus, therapeutics which prevent T cell inhibition by MDSCs should be used to relieve the tumor burden. These should be used in combination with immunotherapies targeting T cell inhibition by PD-L1-PD-1 (*ρ*), such as anti-PD-1 or anti-PD-L1, since varying *ρ* also minimized the tumor size (Fig. 2).

## 4 Parameter Analysis

In this section, we analyze the parameter space and its effect on system (1) by implementing the Approximate Bayesian Computation (ABC) rejection method (Section 4.2) to compare with experimental data (Section 4.1), plotting numerical simulations (Section 4.3), and performing global sensitivity analysis using the extended Fourier Analysis Sensitivity Test (eFAST) (Section 4.4). Section 4.5 contains an examination of the results.

### 4.1 Experimental Data

The model of GBM–immune dynamics (1) developed in this study is applied to a set of experimental data from a high-grade murine glioma cell line, KR158.

Mice were anesthetized and surgically implanted with 35,000 glioma cells through an incision in the skull. At several time points (7, 13, 20, 24, 27, and 34 days after glioma cell implantation), 1–4 mice were euthanized followed by resection of the brain. 2D images of the glioma tissue were used to determine tumor, T cell, and MDSC cell counts (Fig. 3 and Online Resource 2).

**Fig. 3:**
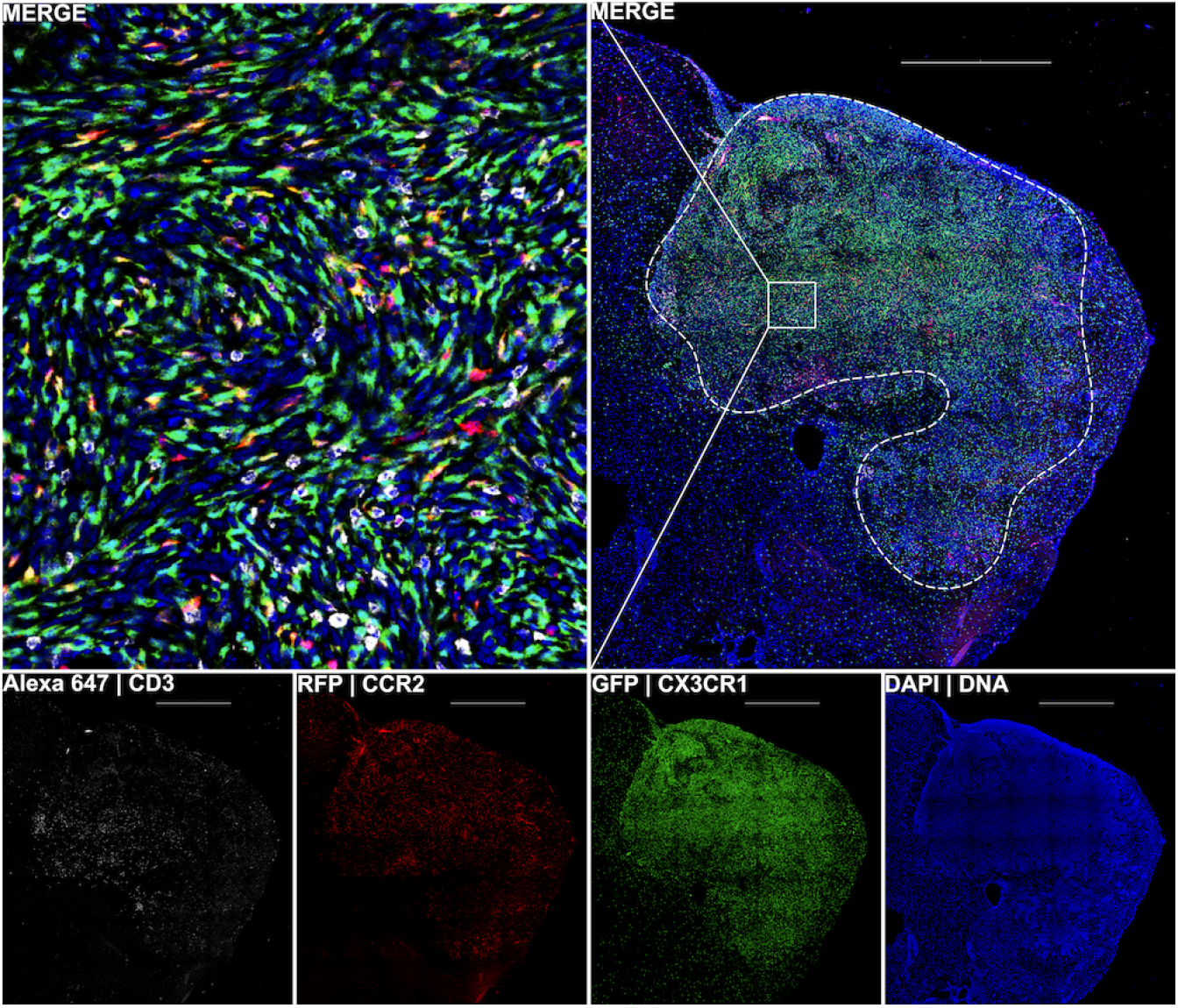
Fluorescent images of brain tissue in a glioma-bearing mouse at day 34 after implantation. CD3 is a marker for T cells, while MDSCs are identified by the CCR2 and CXC3R1 markers. The white, red, and green fluorescent labels denote CD3^+^, CCR2^+^, and CX3CR1^+^ cells, respectively, while the blue fluorescent label denotes all viable cells. The bottom four images are overlaid in the top right image, which is then magnified (top left). A dotted white line marks the glioma’s location. Notice that MDSCs are largely absent from surrounding normal brain tissue.

For a more detailed description of the experimental setup, image analysis pipeline, and data conversion from a 2D section to a 3D spherical tumor, see Online Resource 1 (Secs. 1 and 2).

### 4.2 Approximate Bayesian Computation

Deterministic approaches, like the gradient descent method, are commonly used to estimate a single set of parameter values which optimally represent the given data (Allmaras et al (2013)). This optimization is computed by minimizing error between the numerical simulation and data points. However, depending on the initial guesses of these parameters, this method can converge to different local error minima, making it difficult to uncover a set of parameter estimates which is globally optimal. Additionally, these approaches do not quantify uncertainty in the parameter estimation and generally may not give reliable fits for small data sets. To remedy these issues, we utilize the Approximate Bayesian Computation (ABC) rejection method (Sunnåker et al (2013), Liepe et al (2014)). This algorithm produces probability distributions of the parameter values, which can indicate a range of near-optimal values for each parameter. These distributions provide more detailed information should additional data be made available for a more precise data fitting. In recent years, the use of the ABC rejection method in the study of complex biological systems has been increasing (Browning et al (2017), Stepien et al (2019), Xiao et al (2021), da Costa et al (2018)).

The first step of the ABC rejection method is to specify a prior distribution for each of the parameters. Consequently, we generate 1 million parameter sets, i.e., 1 million 13-dimensional vectors, where each vector forms a parameter set

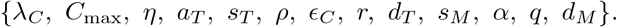

The values for each parameter were chosen uniformly from the ranges shown as the bounds of the horizontal axes in Fig. 4.

**Fig. 4:**
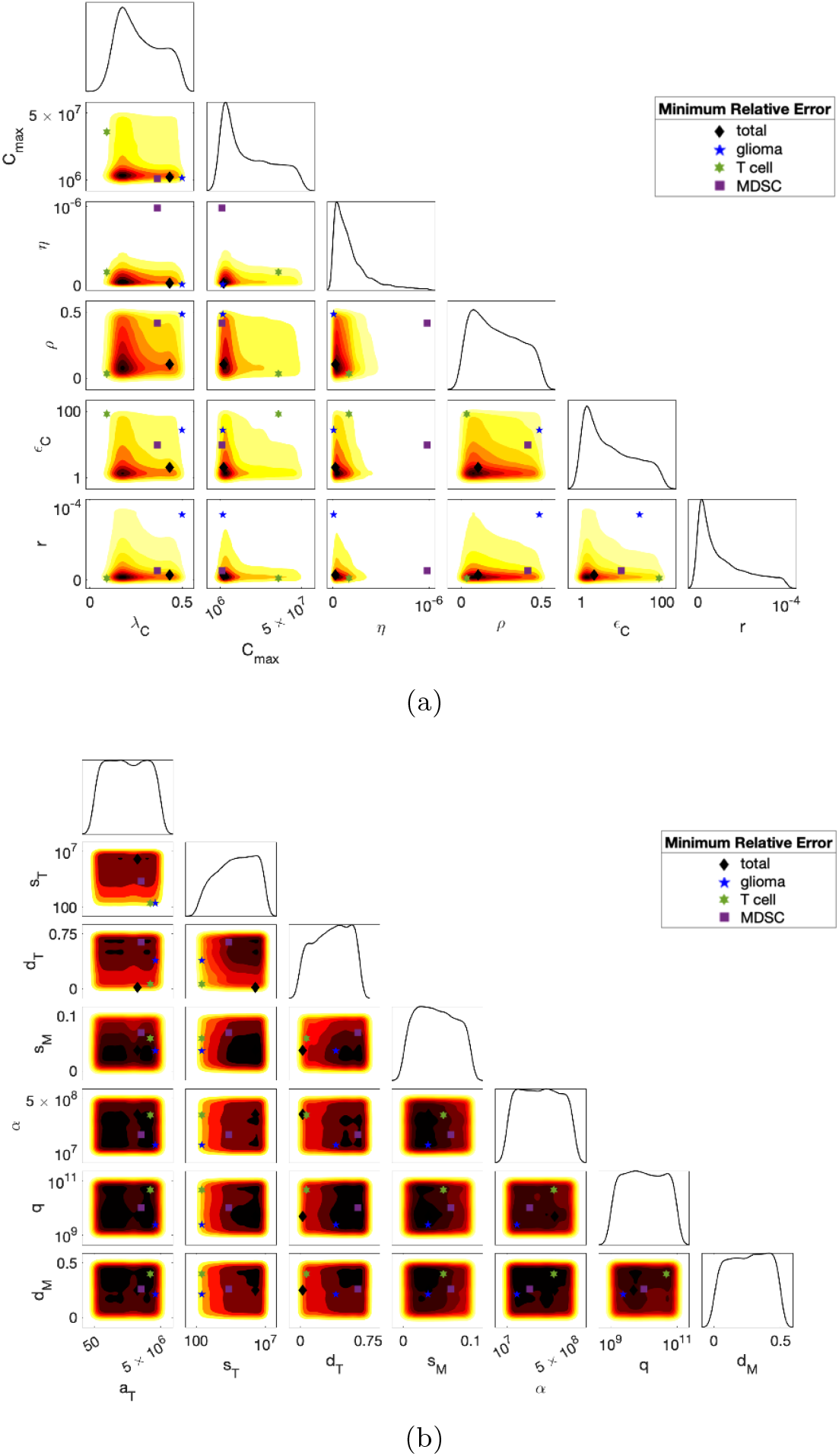
ABC rejection method: The smoothed histograms for the 1D projections of the posterior distributions of each parameter along the diagonal and the 2D contour plot projections for pairs of parameters below the diagonal were produced using the smallest 25% (in terms of the total relative error *E*_total_ (28)) of the accepted parameter values. Darker red indicates a higher frequency of parameter values. The markers indicate the parameter sets with the smallest total relative error (*E*_total_), tumor cell relative error (*E*_*C*_), T cell relative error (*E*_*T*_), and MDSC relative error (*EM*).

Next, we calculate the relative error of a single parameter set,

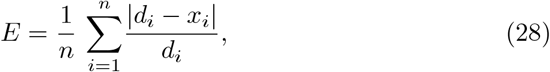

where *d*_*i*_ is the experimental data point and *x*_*i*_ is the simulated value at the *i*^th^ time point. We set *d*_*i*_ to be the average cell count for the *i*^th^ time point, where the time points were days 7, 13, 20, 24, 27, and 34. The relative errors for tumor cells (*E*_*C*_), T cells (*E*_*T*_), and MDSCs (*E*_*M*_) are calculated individually and the total error is *E*_total_ = *E*_*C*_ + *E*_*T*_ + *E*_*M*_ .

Lastly, we specify an error threshold, *R*: if the error of a parameter set is less than the threshold, the set is accepted (i.e., the simulation is sufficiently close to the experimental data), but if it is greater than the threshold, the parameter set is rejected. Since we observed that parameter sets that matched the tumor cell and MDSC data well often did not match the T cell data, we set error thresholds separately for each cell type. In particular, we set *R*_*C*_ = 0.75 for the tumor cell error threshold, *R*_*T*_ = 0.72 for the T cells, and *R*_*M*_ = 0.78 for the MDSCs. For a parameter set to be accepted, we required *E*_*C*_ ≤ *R*_*C*_, *E*_*T*_ ≤ *R*_*T*_, and *E*_*M*_ ≤ *R*_*M*_ .

Of the 1 million parameter sets sampled, approximately 44,000 sets were accepted. Visualizations of the resulting posterior distributions are shown in Fig. 4. Along the diagonal of Fig. 4a are smoothed 1D histograms for parameters exhibiting a right-skewed distribution, while Fig. 4b shows the remaining parameters. Below the diagonal of both of these figures are 2D projections of pairs of parameters. Red shaded areas correspond with higher frequency of the parameter in the posterior distribution.

Table 2 lists the summary statistics of the posterior distributions as well as the best fitting probability distribution, which was calculated by minimizing the 1-Wasserstein metric (earth mover’s distance).

**Table 2:** Parameter summary statistics obtained from the ABC rejection method (Fig. 4) using the smallest 25% (in terms of the total relative error *E*_total_ (28)) of the accepted parameters. Probability distributions which best fit the posterior distributions in Fig. 4 were calculated by minimizing the 1-Wasserstein metric (earth mover’s distance).

|  | Mean | Median | Mode | Standard<br>Deviation | Distribution |
| --- | --- | --- | --- | --- | --- |
| **Glioma Cells** |  |  |  |  |  |
| $\lambda_C$ | 0.272 | 0.250 | 0.174 | 0.114 | Gamma(5.40, 0.050) |
| $C_{\text{max}}$ | $1.87 \times 10^7$ | $1.45 \times 10^7$ | $4.19 \times 10^6$ | $1.48 \times 10^7$ | Logistic($1.74 \times 10^7$, $8.89 \times 10^6$) |
| $\eta$ | $1.84 \times 10^{-7}$ | $1.27 \times 10^{-7}$ | $4.03 \times 10^{-8}$ | $1.86 \times 10^{-7}$ | Weibull($1.84 \times 10^{-7}$, 0.999) |
| **T Cells** |  |  |  |  |  |
| $a_T$ | $2.48 \times 10^6$ | $2.45 \times 10^6$ | $1.98 \times 10^6$ | $1.45 \times 10^6$ | Logistic($2.48 \times 10^6$, $8.80 \times 10^5$) |
| $s_T$ | $5.73 \times 10^6$ | $5.91 \times 10^6$ | $8.79 \times 10^6$ | $2.63 \times 10^6$ | Uniform($10^2$, $10^7$) |
| $\rho$ | 0.223 | 0.207 | 0.0768 | 0.142 | Exponential(0.223) |
| $\epsilon_C$ | 37.5 | 31.1 | 7.00 | 29.3 | Exponential(37.5) |
| $r$ | $2.85 \times 10^{-5}$ | $1.83 \times 10^{-5}$ | $3.60 \times 10^{-6}$ | $2.76 \times 10^{-5}$ | Gamma(0.801, $3.56 \times 10^{-5}$) |
| $d_T$ | 0.402 | 0.415 | 0.656 | 0.213 | Weibull(0.447, 1.81) |
| **Myeloid-Derived Suppressor Cells** |  |  |  |  |  |
| $s_M$ | 0.0492 | 0.0480 | 0.0249 | 0.0278 | Uniform(0, 0.1) |
| $\alpha$ | $2.52 \times 10^8$ | $2.53 \times 10^8$ | $8.27 \times 10^7$ | $1.42 \times 10^8$ | Logistic($2.52 \times 10^8$, $7.82 \times 10^7$) |
| $q$ | $5.09 \times 10^{10}$ | $5.03 \times 10^{10}$ | $3.89 \times 10^{10}$ | $2.83 \times 10^{10}$ | Uniform($10^9$, $10^{11}$) |
| $d_M$ | 0.258 | 0.263 | 0.419 | 0.143 | Normal(0.258, 0.143) |

To determine whether our results were robust, we extracted different percentages of the accepted parameters and compared parameter distributions. Distributions remained largely unchanged when we analyzed any percentage of our accepted parameters.

Since the posterior distributions for parameters in Fig. 4b have less distinct peaks (cf. Fig. 4a), this suggests that, given our data, we were unable to attain additional information on the actual probability distributions for these parameters beyond upper and lower bounds found in the literature.

### 4.3 Numerical Simulations

Since mice were orthotopically implanted with 35,000 KR158 glioma cells in the experiments (Section 4.1), myeloid-derived suppressor cells (MDSCs) are absent from the brain before tumor introduction, and the number of activated T cells are initially negligible, the initial conditions for numerical simulations were set to be *C*(0) = 35, 000, *T* (0) = 0, and *M* (0) = 0 cells. We assume that any loss of glioma cells due to implantation is negligible.

We ran 10,000 simulations using parameter sets sampled from the ABC method posterior distributions listed in Table 2, and illustrate the mean cell counts and cell counts within standard deviation *σ/*4, *σ/*2, 3*σ/*4, and *σ* calculated at each hour in Fig. 5a. We additionally plot the experimental data points for glioma cells, T cells, and MDSCs and observe that the simulations capture nearly all data points within one standard deviation from the mean.

**Fig. 5:**
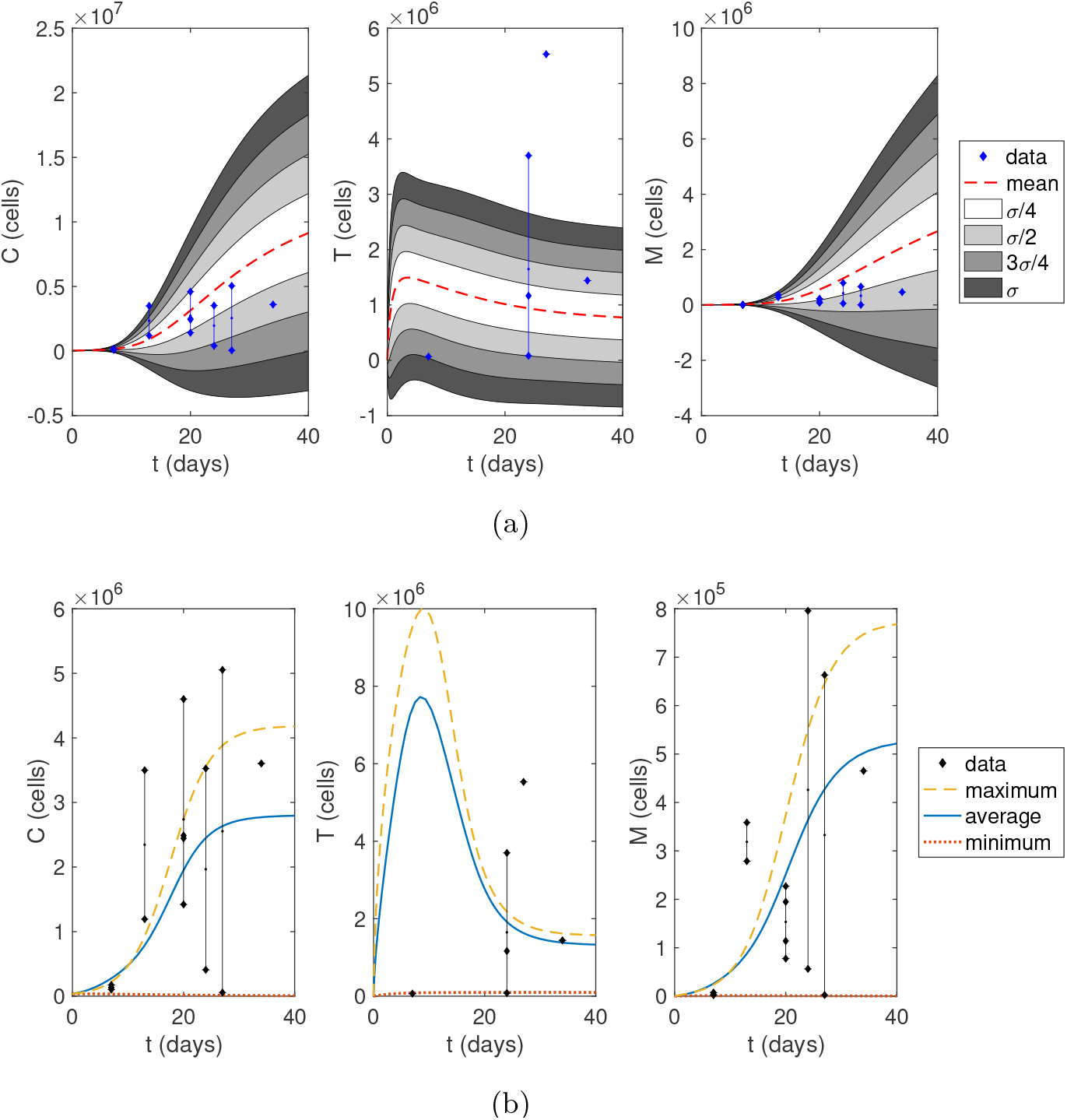
Numerical simulations after implanting 35,000 glioma cells. (a) We sampled 10,000 parameters from the ABC method posterior distributions listed in Table 2. Then, we calculated the mean cell counts at each hour and standard deviation. The simulations capture nearly all data points within one standard deviation from the mean. (b) The maximum, average, and minimum curves represent simulations using the parameter sets of minimal error found using the ABC rejection method (Table S1 (Online Resource 1, Sec. 4)) with respect to the maximum, average, and minimum data points at each time step.

The ABC rejection method histograms illustrated in Fig. 4 were produced by calculating the relative error (28), where *d*_*i*_ are the cell counts averaged at each time point. However, we also used the minimum and maximum cell counts for *d*_*i*_ to evaluate the relative error in two separate instances of the ABC method. The parameter sets that gave rise to minimal error from these three different trials are listed in Table S1 (Online Resource 1, Sec. 4) and numerical simulations are shown in Fig. 5b. A comparison of their total error values could suggest that our parameter ranges are better suited for larger gliomas (error: 0.937) rather than smaller gliomas (error: 1.78), while they are best suited for the average glioma size (error: 0.785). However, these differences in error could also be explained by the variability in the data.

We find that the parameter sets from these three trials (Table S1 in the Online Resource 1, Sec. 4) each satisfy the conditions for a saddle tumor-free equilibrium (Theorem 4), thus corresponding to the worst treatment scenario. We cannot guarantee the existence of a tumorous equilibrium, (*C*^∗^, *T*^∗^, *M*^∗^), for the minimum and average data parameter sets since they fail to fulfill *C*_max_*ϵ*_*C*_*η/λ*_*C*_ < 1 in Theorem 6 given in Appendix A. However, the maximum data parameter set exhibits a tumorous equilibrium when *T*^∗^ < 5.92 × 10^8^ = *λ*_*C*_*/η*. Further, if this equilibrium exists, it is guaranteed to be unique and it is locally asymptotically stable when *T*^∗^ ∈ (11, 2.96 10^8^). All T cell data points are contained within this range, so it is likely that large tumors have a unique, locally asymptotically stable tumorous equilibrium, making these tumors difficult to treat as the system tends towards a tumorous state.

### 4.4 Sensitivity Analysis

To determine the influence of parameters on tumor progression, we conducted a sensitivity analysis using the extended Fourier Analysis Sensitivity Test (eFAST) method outlined in Saltelli et al (1999) by utilizing the MATLAB code developed by the Kirshner Lab at the University of Michigan (Kirschner and Panetta (1998); Kirschner (2008)). eFAST is a global sensitivity analysis method which neither requires model linearity nor a monotonic relationship between the output and the parameters (Saltelli et al (2008)). Similar to the Sobol’ method, it is a variance-based method, but it uses a monodimensional Fourier expansion of the model to evaluate variance. Because of this monodimensional transformation, each parameter *i* can be sampled within the unit hypercube by varying *s* ∈ (− *π, π*) along the space-filling search curve, *x*_*i*_, defined by

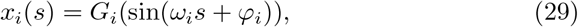

where *ω*_*i*_ is the angular frequency assigned to parameter *i, φ*_*i*_ is a random phase-shift, and *G*_*i*_ is a transformation function.

The ABC posterior distributions of each parameter (Fig. 4 and Table 2) are used to determine the transformation function, *G*_*i*_, hence the search curve, *x*_*i*_. For parameters with right-skewed posterior distributions (Fig. 4a: *λ*_*C*_, *C*_max_, *η, ρ, ϵ*_*C*_, *r*), we used the search curve proposed by Cukier et al (1973),

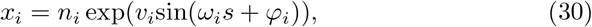

where *n*_*i*_ is the nominal value of the *i*^th^ parameter, set to be the mode of the ABC posterior distribution, and *v*_*i*_ accounts for uncertainty by altering the width of the parameter range. *v*_*i*_ was set to be such that parameters are sampled within the interval [0, 1] by setting *v*_*i*_ = ln(1*/n*_*i*_).

For the remaining parameters (Fig. 4b: *a*_*T*_, *s*_*T*_, *d*_*T*_, *s*_*M*_, *α, q, d*_*M*_), we used the search curve for a uniform distribution that was determined by Saltelli et al (1999),

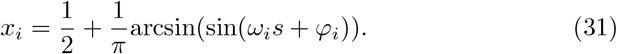

Using these search curves, we generated 2,049 parameter sets for each 5 resamplings, which resulted in a total sample size of 10,245 parameter sets.

Details on how we chose the sample size and number of resamplings are in Online Resource 1 (Sec. 6.1). After parameters were sampled from the unit hypercube, the samples for each parameter were translated to the ranges shown as the bounds of the horizontal axes in Fig. 4.

The main effect, or first-order index, *S*_*i*_, describes the effect that a single parameter *i* has on the variance of the system independently of other parameters. In contrast, the total effect index, *S*_*Ti*_, is a sum of *S*_*i*_ and the higher-order interactions of parameter *i* with other parameters. In Fig. 6, we illustrate the main effect *S*_*i*_ and the total effect index *S*_*Ti*_ for each parameter calculated after a simulation time of 40 days (the approximate time at which an untreated mouse requires euthanasia). The larger the difference between *S*_*Ti*_ and *S*_*i*_, the more interaction there is between *i* and other parameters in synergistically influencing the variance of the output.

**Fig. 6:**
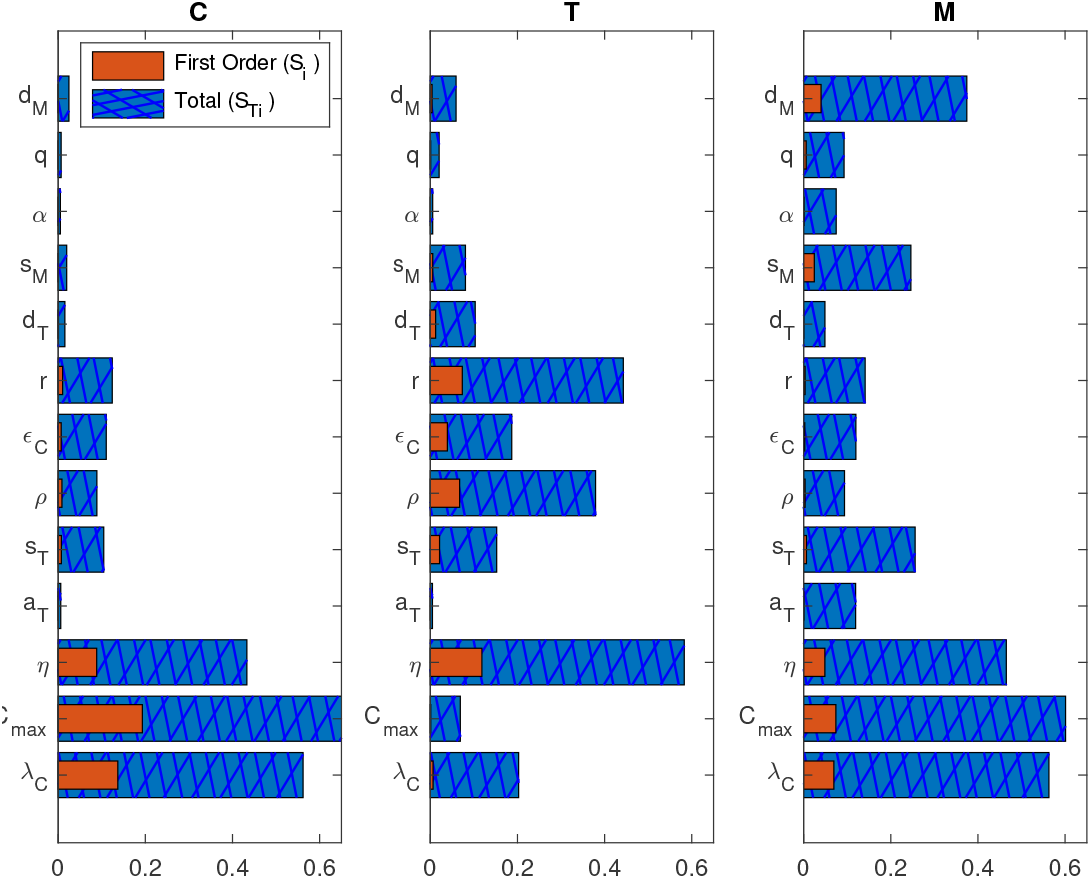
Global sensitivity analysis: The eFAST method was used to calculate the main effect, *Si*, and total effect, *ST i*, of each parameter on tumor cells (*C*), T cells (*T*), and MDSCs (*M*) at *t* = 40 days. Parameters *λC, C*_max,_ *η, ρ, ϵC*, and *r* were sampled using (30), while the remaining were sampled uniformly using (31).

Fig. 6 indicates that T cells are sensitive to perturbations in several parameters, including the kill rate of tumor cells by T cells (*η*) as well as the inhibition of T cells by PD-L1-PD-1 (*ρ*) and MDSCs (*r*). Tumor cells express sensitivity to the tumor growth rate (*λ*_*C*_), tumor carrying capacity (*C*_max_), and the kill rate by T cells (*η*). These results are similar for the MDSC population, however, the death rate (*d*_*M*_) holds a significantly larger influence on variations. Also, while MDSCs and tumor cells display similar total order indices for *η* and *λ*_*C*_, the first order indices of these parameters are larger for tumor cells than MDSCs. This is most likely since these parameters directly influence the tumor cell population but indirectly affect the MDSC population.

Additional sensitivity analyses were run with simulation ending times of 5, 10, 20, 60, 80, and 100 days. Results for days 60, 80, and 100 were qualitatively similar to Fig. 6, and results for days 5, 10, and 20 are shown in Fig. S2 in Online Resource 1 (Sec. 6.2). Comparing the outputs, we find that each cell population becomes increasingly sensitive to *C*_max_ as time progresses, as limited space and nutrients become more significant for larger tumors. MDSCs are initially very sensitive to the MDSC recruitment rate (*s*_*M*_), but this decreased with time and, by day 40, its influence is surpassed by *d*_*M*_ . This result suggests that a CCR2 antagonist should be used as soon as possible, and if the tumor is in its later stages, therapeutics that kill MDSCs at the tumor site should be considered instead. T cells initially were the most sensitive to *ρ* but this switched to *η* by day 10. Further, T cell sensitivity to *r* continues to increase until it surpasses *ρ* by day 20, where it remains the second most influential parameter. Therefore, while treatment with an immune checkpoint inhibitor, such as anti-PD-1 or anti-PD-L1, should be beneficial at any time, results indicate that it is best to treat sooner if possible and to incorporate treatment strategies targeting T cell inhibition by MDSCs later in treatment.

Since the search curve (30) of Cukier et al (1973) produces a stronger skew than our ABC distributions suggest (Fig. 4), we additionally sampled all parameters uniformly using (31) (Fig. S3 in Online Resource 1, Sec. 6.2). We assume that the actual sensitivity indices are within the range produced by the results in Figs. 6 and S3. Fig. S3 shows slightly more interaction occurring between parameters (as seen by greater differences between *S*_*i*_ and *S*_*Ti*_), thus, by utilizing the ABC posterior distributions to inform the search curve, Fig. 6 more precisely determined which parameters should be targeted with treatment.

### 4.5 Results

In the existing literature, we could find no estimates for the value of *s*_*M*_ . From the ABC posterior distribution, we expect *s*_*M*_ to fall within the range of 0.0214–0.0770 day^−1^ (Table 2). Separately, we approximated *s*_*M*_ using data on CCL2 expression in KR158 glioma cells as well as the migratory effect of CCL2 on MDSCs in vitro (Online Resource 1, Sec. 3.1) and found that KR158 gliomas recruit MDSCs at a rate of 0.0123–0.0144 MDSCs per day per tumor cell. Since this data only included the effect of CCL2 and not CCL7 on MDSC migration, we expect this range to be an underestimate for *s*_*M*_ . However, as the mode of the *s*_*M*_ distribution, 0.0249, is close to double the experimentally calculated value, we expect that CCL7 will have a similar effect on MDSC migration as CCL2, assuming that the interaction between CCL2 and CCL7 causes an additive effect in recruitment.

In all three populations, the tumor growth rate (*λ*_*C*_), tumor carrying capacity (*C*_max_), and the tumor kill rate by T cells (*η*) are sensitive parameters which exhibit interactions with other parameters, as seen by large differences between the first and total order indices. Although the sensitivity analysis indicates this interaction (Fig. 6), it does not identify which parameters are interacting with each other. By combining the sensitivity analysis results with the ABC histograms in Fig. 4a, we can conjecture potential relationships.

There is a considerable inverse relationship between *λ*_*C*_ and the PD-L1-PD-1 parameters (*ρ* and *ϵ*_*C*_), *C*_max_ and the inhibition by PD-L1-PD-1 (*ρ* and *ϵ*_*C*_) and MDSCs (*r*), as well as *η* and *r* and slightly between *η* and *ϵ*_*C*_. Although these inverse relationships could potentially be due to these parameters each exhibiting right-skewed distributions, the inverse relationships between *λ*_*C*_, *C*_max_, and *η* themselves are negligible. Therefore, we hypothesize that, although the tumor population is less sensitive to the immune suppression parameters *ρ, ϵ*_*C*_, and *r*, targeting these would affect the tumor’s sensitivity to *λ*_*C*_, *C*_max_, and *η*. At the very least, according to the T cell sensitivity in Fig. 6, targeting these parameters would increase the immune response, thus decreasing the tumor population.

Fig. 4a also displays an inverse relationship between the tumor upregulation of PD-L1 (*ϵ*_*C*_) and the inhibition of T cells by the PD-L1-PD-1 complex (*ρ*) and by MDSCs (*r*). The relationship between *ρ* and *ϵ*_*C*_ suggests that as tumor cells express more PD-L1, the PD-L1-PD-1 complex need not be as effective at inhibition in order to produce the same outcome. The second relationship suggests that *r*, although it is a less sensitive parameter, helps to increase the sensitivity of the system to *ϵ*_*C*_. Thus, perturbations in the two forms of immunosuppression, the PD-L1-PD-1 complex and MDSCs, cause greater variance on the tumor together rather than alone.

## 5 Discussion and Future Directions

Current treatment paradigms for the highly aggressive brain tumor glioblastoma (GBM) present limited efficacy. Barriers, such as the highly immune suppressed tumor microenvironment, make gliomas difficult to treat. To address this barrier, we mathematically model GBM-immune dynamics by considering the role of myeloid-derived suppressor cells (MDSCs) in aiding GBM progression through T cell suppression. Gliomas recruit CCR2^+^ MDSCs to the tumor microenvironment by expressing the chemokines CCL2 and CCL7, which are ligands of the CCR2 receptor. Once at the tumor site, MDSCs suppress T cells, which are already inhibited by the formation of the PD-L1-PD-1 complex. This complex forms due to T cell interaction with tumor cells, and it masks the tumor from identification by T cells. Incorporating these two forms of immunosuppression prepares our model for future extension to include immunotherapies and optimize treatments.

Our results reinforce pre-clinical studies by Flores-Toro et al (2020) which suggest that MDSCs and the PD-L1-PD-1 immune checkpoint should be targeted together to increase the immune response and thus decrease the tumor burden. Therefore, promising therapeutics include immune checkpoint inhibitors, such as anti-PD-1 and anti-PD-L1, and CCR2 antagonists. Time-dependent sensitivity analysis with eFAST indicates that treatment with these therapeutics should be used as soon as possible; however, an immune check-point inhibitor should offer some benefit at any time. In contrast, MDSC recruitment should be blocked by a CCR2 antagonist in the early stages of a tumor, but immunotherapies which kill MDSCs at the tumor site should be considered for larger tumors. While CCR2 antagonists prevent MDSC recruitment (thus targeting the *s*_*M*_ parameter), strategies which directly target the parameter *r* by decreasing the ability of MDSCs to inhibit T cells should be explored. Time-dependent eFAST results suggest that therapies targeting *r* would be especially useful during the later stages of tumor development.

Stability analysis of the system (1) established conditions for the existence and stability of the tumor-free equilibrium, (0, *T*_0_^∗^, 0). This unique equilibrium is corroborated by immunohistochemistry data showing the absence of MDSCs within the tumor-free mouse brain (Fig. 3). When 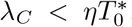, i.e., the initial immune response is greater than the tumor growth rate, the tumor-free equilibrium is locally asymptotically stable, which is equivalent to results in Nikolopoulou et al (2018). Further, it is globally stable when 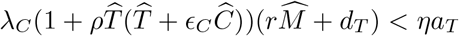 . This suggests that, regardless of the initial number of activated T cells, if T cells are activated and kill tumor cells faster than tumor cells proliferate and faster than T cells are suppressed and die naturally, then the system will tend to the tumor-free equilibrium.

Bifurcation analysis further illuminates this result by showing that shifting only 4 parameters (*λ*_*C*_, *η, ρ, r*) can result in the system tending to the tumor-free equilibrium. As surgical resection of a tumor can decrease the tumor growth rate by disrupting the influx of nutrients through damage to the vasculature, these results suggest that a treatment plan including surgery and immunotherapies which target the PD-L1-PD-1 complex and the immunosuppressive capabilities of MDSCs would minimize the tumor size.

We obtained estimates of the tumor volume and cell counts from murine GBM data collected from immunofluorescence imaging. T cell data was more sparse than the glioma and MDSC data, so future model predictions could be improved by incorporating more time-dependent T cell data. Numerical simulations in Fig. 5a show that our parameter distributions capture nearly all data points (33 of 36 data points) within one standard deviation from the mean and most (27 of 36) within half of a standard derivation. In general, more data would improve the predictive capabilities of the model, especially if the cell populations within a single mouse could be tracked over time.

Given the limited data set, we used the Approximate Bayesian Computation (ABC) rejection method in unison with data to obtain information on the probability distribution of realistic parameter values. We identified approximate values and quantified the associated uncertainty for previously unknown parameters, in particular, the recruitment rate of MDSCs (*s*_*M*_) and the inhibition rate of T cells by MDSCs (*r*). By calculating an estimate of *s*_*M*_ using chemokine expression levels on glioma cells and MDSC migration in response to certain concentrations of the chemokine CCL2 measured from experimental data, we found that the parameter values estimated by the ABC method were of the same order as calculated from data. Further, we found that the values estimated for *r* were noticeably greater than the estimated values for *η* (Table 2), suggesting that MDSCs suppress T cells more effectively than T cells kill tumor cells. This would explain the difficulty T cells experience in overcoming a glioma. Further biological testing would need to occur to validate this hypothesis.

The ABC parameter distributions (Fig. 4) informed global sensitivity analysis by identifying a search curve with which to sample parameters for eFAST. Sensitivity analysis (Fig. 6) indicated that there are several drivers of the system, including the tumor growth rate (*λ*_*C*_), tumor carrying capacity (*C*_max_), and T cell kill rate (*η*), along with noticeable interaction occurring between parameters to affect the variance of the system. By comparing the eFAST results with the ABC results, we hypothesize that interaction could be occurring between these three drivers and parameters representing immune suppression–namely the tumor upregulation of PD-L1 (*ϵ*_*C*_) along with the inhibition by PD-L1-PD-1 (*ρ*) and by MDSCs (*r*), since there appears to be an inverse relationship between these parameters in Fig. 4a. Further, T cell sensitivity in Fig. 6 shows that targeting these parameters would increase the immune response regardless of any interactions. This suggests that the PD-L1-PD-1 complex and inhibition by MDSCs should be therapeutically targeted together to relieve tumor burden and increase the immune response.

Overall, there is a mutually beneficial relationship between the ABC method and eFAST. The distributions produced by ABC in accordance with data inform the choice of search curve when sampling parameters in eFAST. In return, the interaction effects displayed by eFAST direct us to look for relationships between parameters using the ABC results. Thus, the two together increase the reliability of results as well as the inferences that can be made.

Future directions include extending this model to incorporate immunotherapies targeting the PD-L1-PD-1 complex and MDSCs and then optimizing treatment regimens to minimize tumor burden. Anti-PD-1, which targets PD-1 on T cells, has not been successful as a monotherapy in part because it requires more activated T cells to be in the tumor microenvironment (Kleponis et al (2015)). Simulations in Nikolopoulou et al (2018) suggest that anti-PD-1 alone is not able to increase the number of T cells to a tumor eradication threshold. Through inhibition, MDSCs decrease the number of activated T cells. Treatment with a CCR2 antagonist decreases the recruitment of MDSCs, which indirectly increases the number of activated T cells, thus improving the efficacy of anti-PD-1. This improved efficacy is supported by data showing increased survival rates in glioma-bearing mice treated with anti-PD-1 and the CCR2 antagonist, CCX872 (Flores-Toro et al, 2020, Fig. 4). While our results agree with this outcome, we also propose an additional target. Collectively, our global stability and bifurcation analyses, sensitivity analysis, and ABC method results suggest that it would be more beneficial to directly target the mechanisms by which MDSCs inhibit T cells rather than MDSC recruitment. Therefore, therapeutics which prevent MDSC inhibition of T cells should be tested in combination with immune checkpoint inhibitors like anti-PD-1.

## Supporting information

Online Resource 1

## Statements and Declarations

## Acknowledgments

The authors would like to thank and acknowledge the reviewers for the time, effort, expertise, and comments given on the original manuscript.

## Ethics approval

All procedures involving animal housing and surgical protocols were followed according to the guidelines of the University of Florida Institutional Animal Care and Use Committee.

## Authors’ contributions

Conceptualization and Methodology: all authors; Formal analysis, Software, Validation, and Visualization: Hannah G. Anderson; Investigation and Data Curation: Gregory P. Takacs; Resources: Jeffrey K. Harrison; Supervision: Tracy L. Stepien, Jeffrey K. Harrison, Yang Kuang; Writing - original draft preparation: Hannah G. Anderson; Writing - review and editing: all authors.

## Code availability

The source code used to generate the results for this article is available through GitHub at https://github.com/stepien-lab/glioma-Tcell-MDSC [v1.0.0]. The code is platform independent and written in MATLAB.

## Supplementary information. Online Resource 1

*ESM1.pdf* contains details regarding data collection and usage, the literature search and estimation of parameters, as well as additional figures and information about the ABC rejection method, bifurcation diagrams, and eFAST.

### Online Resource 2

*ESM2.xlsx* contains time-dependent experimental data of tumor volumes and cell counts of the three studied cell populations, which were collected according to the methods described in Online Resource 1.

## Appendix A

### Additional Theorems

In the following appendix, we establish conditions for the existence and stability of a unique tumorous equilibrium.

#### Theorem 6

(Uniqueness of tumorous equilibrium) *The system* (1) *has a tumorous equilibrium if λ*_*C*_ *> ηT*^∗^ *and C*_max_*ϵ*_*C*_ *η/λ*_*C*_ < 1. *Further, it is unique when ϵ*_*C*_ *ρ > η/λ*_*C*_, *C*_max_ ≥ 2, *β*_1_*/β*_2_ < *C*_max_*ϵ*_*C*_ *λ*_*C*_ */η, and ρβ*_1_ (*λ*_*C*_ */η*)^2^ < *C*_max_ (*s*_*M*_ *r* − *d*_*M*_ *s*_*T*_), *where β*_*i*_ = *d*_*T*_ *d*_*M*_ + *αr* + *s*_*M*_ *r*((*i* + 1)*C*_max_ + *q, for i* = 1, 2.

*Proof* Setting the right hand side of the tumor cell equation (1a) equal to zero and evaluating at the tumorous equilibrium implies that

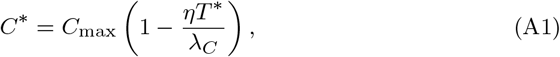

which is positive if

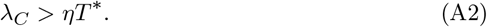

For the MDSCs equation (1c), we find that

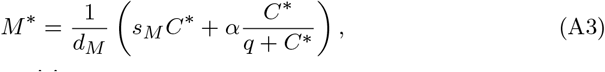

which is positive if *C*^∗^ is positive.

The T cell equation (1b) yields

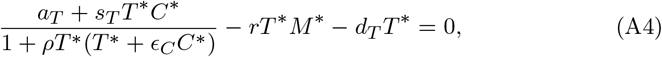

which, when we substitute (A3) and (A1) and rearrange, becomes a degree-5 polynomial,

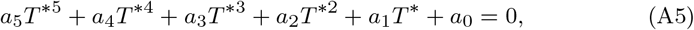

Where

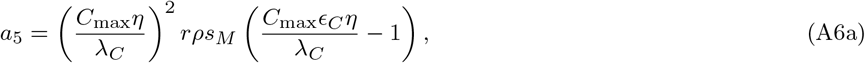

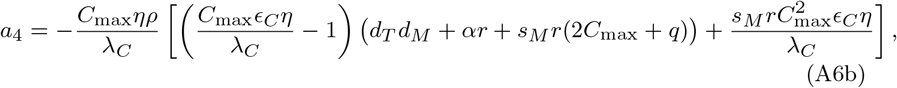

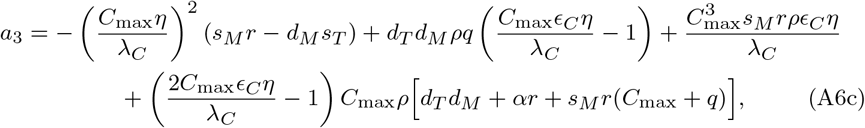

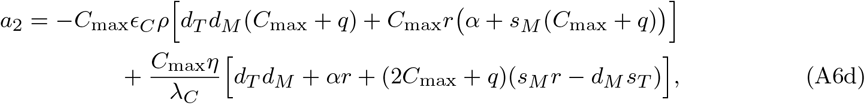

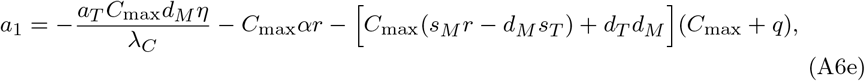

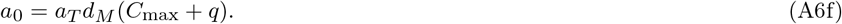

We observe that *a*_0_ is always positive since all the parameters are positive. Since *a*_0_ is positive, when we require *a*_5_ to be negative, the number of sign changes is 1 mod 2 regardless of the signs of the coefficients *a*_1_, …, *a*_4_. Therefore, by Descartes’ rule of signs, there is at least one positive zero for (A5). Thus, a tumorous equilibrium is guaranteed to exist if we enforce condition (A2) and

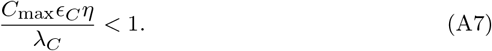

We derive conditions for uniqueness in the case where *a*_*i*_ < 0 for *i* = 1, …, 5. *a*_1_ is negative if we require

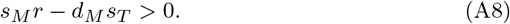

In other words, immunosuppression due to MDSCs must be greater than the recruitment of T cells and death of MDSCs.

For *a*_2_ to be negative, under the conditions that

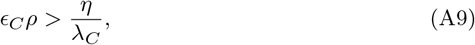

it follows that

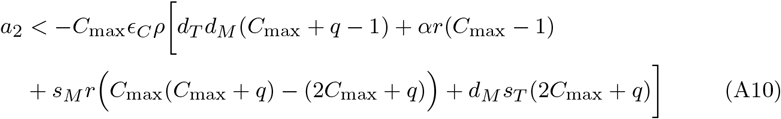

When we require

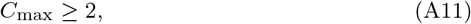

then

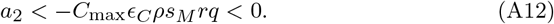

We shall consider *a*_4_ next. When we expand *a*_4_ by distributing the term containing *d*_*T*_ *d*_*M*_, it becomes clearer to see that *a*_4_ is negative under the condition that

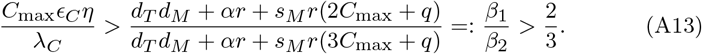

For *a*_3_, conditions (A8) and (A7) cause the first and second terms of *a*_3_ to be negative, respectively, while (A13) implies that the last term is positive. Applying (A7) to the last three terms and simplifying,

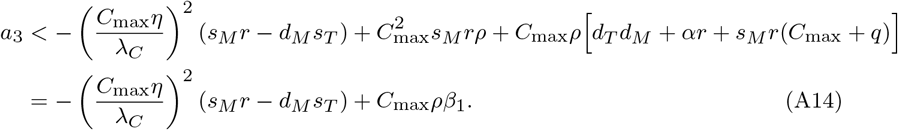

Thus, *a*_3_ is negative under the condition that

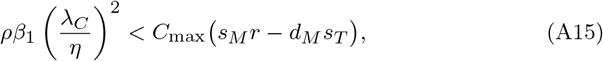

which is a stronger version of (A8).

Under this condition along with (A7), (A9), (A11), and (A13), we have *a*_*i*_ < 0 for *i* = 1, …, 5 and *a*_0_ > 0. Thus, according to Descartes’ rule of signs, there is one unique positive zero for *T*^∗^, and correspondingly from (A1) and (A3), one unique *C*^∗^ and *M*^∗^. Therefore, we have a unique tumorous equilibrium (*C*^∗^, *T*^∗^, *M*^∗^). □

There are some biological implications from these conditions. First of all, condition (A9) can be rewritten as *η < λ*_*C*_*ϵ*_*C*_*ρ*. This suggests that the tumor growth rate (*λ*_*C*_) multiplied by immunosuppression via the PD-L1-PD-1 complex (*ϵ*_*C*_*ρ*) must be larger than the ability of T cells to kill tumor cells (*η*). Further, condition (A15) indicates that *d*_*M*_ *s*_*T*_ < *s*_*M*_ *r*, in other words, immunosuppression due to MDSCs (*s*_*M*_ *r*) must be greater than the recruitment of T cells (*s*_*T*_) and death of MDSCs (*d*_*M*_). Both implications suggest that the suppression of the immune system has to be substantial enough for uniqueness of the tumorous equilibrium to be guaranteed.

Next we determine conditions for the stability of the tumorous equilibrium.

#### Theorem 7

(Stability of tumorous equilibrium) If a tumorous equilibrium, (*C, T, M*) = (*C*^∗^, *T*^∗^, *M*^∗^), exists as determined by the conditions in Theorem 6, it *is locally asymptotically stable when λ*_*C*_ > 2*ηT*^∗^ *and ρT*^∗2^ ≥ 1.

*Proof* The Jacobian evaluated at the tumorous equilibrium is

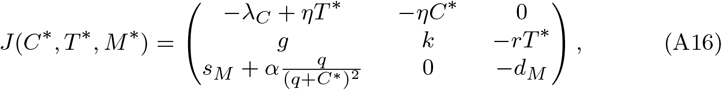

where

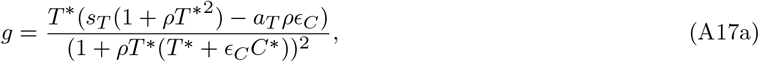

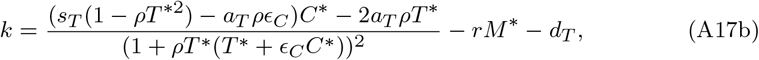

and we have substituted (A1) into the (1, 1)-entry to simplify the expression.

When we require

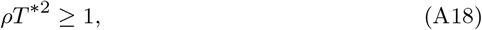

*k* is guaranteed to be negative.

The characteristic polynomial of *J*(*C*^∗^, *T*^∗^, *M*^∗^) is

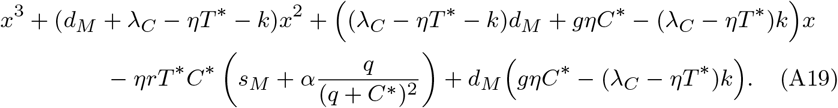

A monic polynomial is defined to be Hurvitz if Re(*x*_*i*_) < 0 for all roots *x*_*i*_ ∈ ℂ. By the Routh-Hurvitz Criterion, a polynomial *p*(*x*) = *x*^3^ + *b*_1_*x*^2^ + *b*_2_*x* + *b*_3_ is Hurvitz if and only if *b*_*i*_ > 0 for *i* = 1, 2, 3, and (*b*_1_*b*_2_ − *b*_3_) > 0.

Under conditions (A2) and (A18) it follows that

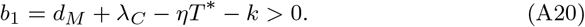

If we strengthen condition (A2) to be

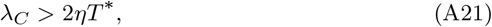

this allows us to conclude, after substituting *g* and *k* and simplifying, that

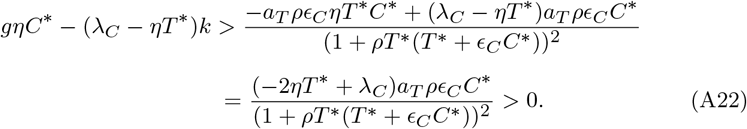

Therefore,

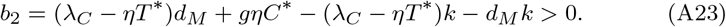

Next, since *rM*^∗^ < −*k*, and substituting in *M*^∗^ (A3), it follows by condition (A21) that

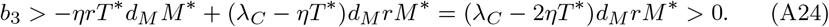

Finally, we shall address the positivity of (b_1_b_2_ − b_3_),

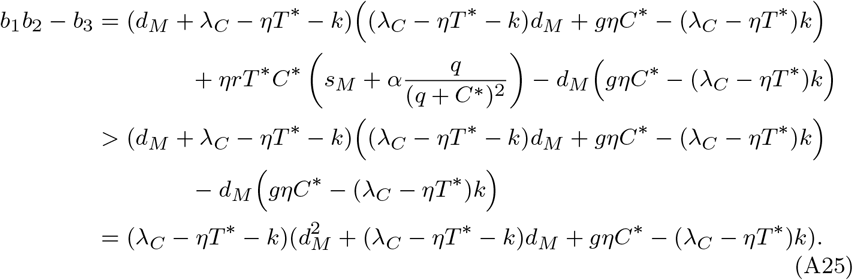

Now, *λ*_*C*_ − *ηT*^∗^ − *k* > 0 since *k <* 0 and by condition (A21). Therefore, by condition (A22),

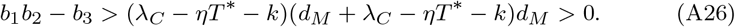

Since *b*_*i*_ > 0 for *i* = 1, 2, 3 and (*b*_1_*b*_2_ −*b*_3_) > 0, the characteristic polynomial (A19) is Hurvitz. Therefore, the tumorous equilibrium is locally asymptotically stable under the conditions (A18) and (A21). □

Condition (A18) requires that the T cell inhibition by the PD-L1-PD-1 complex is at or above a certain level. Condition (A21) effectively assumes that the tumor cell growth rate is larger than twice the kill rate by the T cell population. We note that Nikolopoulou et al (2018) also required a similar condition (*λ*_*C*_ *> ηT*^∗^) in order to achieve a stable tumorous equilibrium for their system.

