## Supplementary material for "Global stability and parameter analysis reinforce therapeutic targets of PD-L1-PD-1 and MDSCs for glioblastoma": Online Resource 1

*Journal of Mathematical Biology*

Hannah G. Anderson[1], Gregory P. Takacs[3], Duane C. Harris[2], Yang Kuang[2], Jeffrey K. Harrison[3] and Tracy L. Stepien[1*]

[1]Department of Mathematics, University of Florida, Gainesville, FL, USA.

[2]School of Mathematical and Statistical Sciences, Arizona State University, Tempe, AZ, USA.

[3]Department of Pharmacology and Therapeutics, University of Florida, Gainesville, FL, USA.

**4 Parameter Sets with Smallest Error** **8**

**5 Additional Bifurcation Diagrams** **9**

**6 Additional eFAST Details and Figures** **10**
6.1 Number of Samples ($\boldsymbol{N_s}$) and Resamplings ($\boldsymbol{N_r}$) . . . . . . . . 10
6.2 Additional Figures . . . . . . . . . . . . . . . . . . . . . . . . . 10

In this online resource, we provide additional appendices to our paper which focused on the following glioblastoma-immune dynamics model:

$$\frac{dC}{dt} = \underbrace{\lambda_C C\left(1 - \frac{C}{C_{\max}}\right)}_{\substack{\text{logistic}\\ \text{growth}}} - \underbrace{\eta T C}_{\substack{\text{killed by}\\ \text{T cells}}}, \tag{1a}$$

$$\frac{dT}{dt} = \Big( \underbrace{a_T}_{\text{activation}} + \underbrace{s_T T C}_{\text{stimulation}} \Big) \underbrace{\frac{1}{1 + \rho T (T + \epsilon_C C)}}_{\substack{\text{inhibition by}\\ \text{PD-1-PD-L1}}} - \underbrace{r T M}_{\substack{\text{inhibition}\\ \text{by MDSCs}}} - \underbrace{d_T T}_{\text{death}}, \tag{1b}$$

$$\frac{dM}{dt} = \underbrace{s_M C}_{\substack{\text{stimulation by}\\ \text{CCL2/CCL7}}} + \underbrace{\alpha \frac{C}{q + C}}_{\substack{\text{expansion}\\ \text{of MDSCs}}} - \underbrace{d_M M}_{\text{death}}. \tag{1c}$$

### 1 Experimental Setup and Image Analysis

The following details the murine glioma experimental study and image analysis pipeline.

#### 1.1 Animals

CX3CR1-deficient CX3CR1$^{\mathrm{GFP/GFP}}$ [B6.129P-Cx3cr1tm1Litt/J]), and CCR2-deficient CCR2$^{\mathrm{RFP/RFP}}$ [B6.129(Cg)-Ccr2tm2.1Ifc/J]) mice were obtained from The Jackson Laboratory. CCR2$^{\mathrm{RFP/WT}}$/CX3CR1$^{\mathrm{GFP/WT}}$ mice were generated via in-house breeding of homozygous CX3CR1$^{\mathrm{GFP/GFP}}$ and homozygous CCR2$^{\mathrm{RFP/RFP}}$ mice. All procedures involving animal housing and surgical protocols were followed according to the guidelines of the University of Florida Institutional Animal Care and Use Committee.

#### 1.2 Cell Culture

KR158B glioma cells were cultured in Dulbecco's Modified Eagle Medium (DMEM) supplemented with 1% penicillin-streptomycin and 10% fetal bovine serum (FBS). Cells were grown in a humidified incubator at 37 °C with 5% CO2. DMEM and penicillin-streptomycin were purchased through Invitrogen. FBS was purchased through Thermo Scientific.

#### 1.3 Orthotopic Brain Tumor Model

Animals were anesthetized using isoflurane and administered analgesia prior to cell injection. While under anesthesia, the surgical site was prepared, a 2- to 3-mm incision was made at the midline of the skull. Using a stereotaxic apparatus (Stoelting), the mice were secured, and a Hamilton syringe was positioned 2-mm lateral from the bregma. KR158B glioma cells ($3.5 \times 10^4$), in a total volume of 2 $\mu$L were injected 3-mm deep into the right cerebral hemisphere using an automated microfluidic injection system (Stoelting) at a rate of 1 $\mu$L/min. Cells are suspended in a 1:1 ratio of methylcellulose to PBS. Post injection, the needle was retracted slowly, and the surgical site was closed via suture and bone wax. Animals were then placed into a warm cage for postsurgical monitoring.

#### 1.4 Immunofluorescence

Mice were euthanized and transcardial perfusions were performed using a 10mL syringe with a 25G winged infusion set, of 20mL 4.0% paraformaldehyde (PFA) solution. Following fixative perfusion, mouse brains were removed and soaked in 4.0% PFA for 1hr. Brains were subsequently transferred to 30% sucrose solution for 24hrs and snap frozen using liquid nitrogen chilled 2-Methylbutane. Brains were embedded in optimal cutting temperature compound and mounted for cryo-sectioning (Lecia Biosystems Cryostat). 5-10$\mu$m thick sections were taken and mounted on microscope slides. Sections were dried overnight at 4C. For CD3$\epsilon$ staining, sections were brought up to room temperature and rehydrate in PBS for 5 mins. Normal goat serum (10%) blocking buffer was applied in a humidified chamber at RT for 1 hr. Rat anti-mouse CD3$\epsilon$ (Biolegend 155602) primary antibody (0.5mg/mL) was applied to each slide and stored at 4C overnight. The following day, samples were washed in PBS 3 times and incubated subsequently in goat anti-rat Alexa 647(Invitrogen A21247) secondary antibody (2 mg/mL). The sections were then washed 3 times with PBS, counterstained with Vectashield antifade mounting medium with 4',6- diamidino-2-phenylindole, and imaged using an inverted Nikon TiE-PFS-A1R confocal microscope. Images were post-processed using Nikon Elements software.

##### 1.5 Cell Quantification

For each brain tumor, 3 sections were imaged using a 20x objective, quantified, and averaged prior to statistical inferences. For larger tumors, a series of 20x images were taken and stitched together using Nikon Elements software. Images were imported into Cell Profiler (Stirling et al (2021)) for quantification using an image-analysis pipeline. Briefly, the pipeline utilizes each fluorescent channel to identify and quantify DAPI-stained nuclei, $CCR2^{WT/RFP}$, $CX3CR1^{WT/GFP}$, and CD3 cells. Masks were generated for each cell identity. DAPI-stained nuclei mask was used as a prerequisite for subsequent cell identification. To identify $CCR2^{WT/RFP}$/$CX3CR1^{WT/GFP}$ cells "RelateObjects" command was carried out to identify co-localization of RFP and GFP positive cells. Tumor area was calculated through the cropping function with dysplastic DAPI staining denoting tumor boundary. The final output includes tumor area and numbers of each cell type.

### 2 Data Usage

The following discusses the data usage as well as statistical analyses performed.

#### 2.1 Data Conversion

Using Cell Profiler (Stirling et al (2021)), the immunofluorescence data measured the number of MDSCs and T cells in 2D on a 0.01 mm thick slice that was removed from the approximate center of the tumor (Appendix 1.5). Since CellProfiler was not able to determine the number of tumor cells in the 2D image, we estimated this by assuming that the density of tumor cells in an epithelial tumor is approximately $10^8$ cell/cm$^3$ (Del Monte (2009)). To convert these cell numbers in terms of a 3D tumor, we assumed that the tumor is spherical and that the average density of cells throughout the tumor is that of the slice. The tumor area on the 2D image was used to find a radius, which was then used to calculate a volume. For each of the cell types, we calculated a 2D density, and then used that along with the volume to calculate the total number of cells throughout the entire 3D tumor.

#### 2.2 Statistical Analysis

We calculated that there were strong outliers for the number of glioma cells and MDSCs measured from two mice in the data by defining a "strong outlier" to be any value which is more than 3 $\times$ IQR above/below the upper/lower quartile, respectively. Although we used these two strong outliers to create an adequate range for the carrying capacity of tumor cells, $C_{\max}$, we eliminated these two data points when running the Approximate Bayesian Computation method (Section 4.2). Since there were less T cell data points, we were unable to show the T cell data from those two mice to be outliers. This is likely why the numerical simulations in Fig. 5 are unable to capture the two largest T cell points within one standard deviation of the mean according to the ABC posterior distributions. Given more data for T cells, these data points might be considered outliers and eliminated.

### 3 Estimated Parameter Values

The following includes information about notable parameter values and the method by which they were estimated for Table 1.

#### 3.1 Glioma Parameter Values

##### Tumor Growth Rate ($\lambda_C$)

Stensjøen et al (2015) calculated growth rates of untreated human glioblastomas in vivo by fitting data to the exponential, linear radial, and Gompertz growth models. The median specific growth rate calculated was 0.014 day$^{-1}$, which we set as the lower bound for $\lambda_C$ in Table 1. Other previous estimates of $\lambda_C$ range from 0.2304–0.264 day$^{-1}$ for in vivo GL261-luc2 cells implanted in rats (Doblas et al (2010)) and 0.4008–0.4512 day$^{-1}$ for the same cell type implanted in mice (Hypothesis 1 in Rutter et al (2017)). Thus, we set 0.4512 day$^{-1}$ as the upper bound for $\lambda_C$ in Table 1.

##### Carrying Capacity ($C_{\max}$)

We calculated the carrying capacity of tumor cells, $C_{\max}$, to have a range of $4.057 \times 10^6$–$4.14 \times 10^7$ cells from the number of glioma cells measured at all time points from the experimental data (Appendix B). The data point $4.14 \times 10^7$ cells was approximately $9.5 \times$ IQR above the upper quartile, so we set this as the upper bound because it likely includes all realistic values for carrying capacity of GBM in mice. The lower bound of $4.057 \times 10^6$ cells was determined by eliminating the two outliers (Appendix B.2) and averaging the five largest tumor cell numbers in the data.

#### 3.2 T Cell Parameter Values

##### Activation Rate ($a_T$)

We derived the activation rate, $a_T$, by combining four parameters from Nikolopoulou et al (2018) into one due to parameter non-identifiability,

$$a_T := \lambda_{TI_{12}} T_N \frac{I_{12}}{K_{I_{12}} + I_{12}} = 2.643 \times 10^{-3} \text{day}^{-1} \text{ g/cm}^3, \tag{2}$$

where $\lambda_{TI_{12}} = 8.81$ day$^{-1}$ is the activation rate of T cells by IL-12, $T_N = 6 \times 10^{-4}$ g/cm$^3$ is the density of naive T cells, $I_{12} = 1.5 \times 10^{-10}$ g/cm$^3$ is the IL-12 concentration, and $K_{I_{12}} = 1.5 \times 10^{-10}$ g/cm$^3$ is the half saturation of IL-12 (Nikolopoulou et al (2018); Lai and Friedman (2017)).

This value of $a_T$ (2) is in units of density, which is in terms of grams of T cells (not IL-12) over volume. Using dimensional analysis, we take into account the range for T cell mass, which Grover et al (2011) measured as 430 pg–1310 pg, along with the minimum and maximum tumor volumes in Online Resource 2 to produce the range for $a_T$ given in Table 1.

Although T cell activation does occur within the glioma microenvironment, it is not necessarily by IL-12, which is implicitly assumed in (2). Nonetheless, the calculation provides a reasonable approximation of the influence of T cell activation on the system.

#### Inhibition of T Cells by Formation of PD-L1-PD-1 Complex ($\rho$)

Similar to $a_T$, we also combined multiple parameters in Nikolopoulou et al (2018) to estimate $\rho$. Nikolopoulou et al (2018) expressed the function for PD-L1-PD-1's suppression of T cells in the treatment-free case by

$$F(C,T) = \frac{1}{1 + \frac{\sigma \rho_P \rho_L}{K_{TQ}} T(T + \epsilon_C C)}, \tag{3}$$

where $\sigma = 1$ cm$^3$/g is the ratio of association to dissociation of the PD-L1-PD-1 complex (assumed to be in equilibrium), $\rho_P = 3.19 \times 10^{-7}$–$8.49 \times 10^{-7}$ is the expression level of PD-1 on T cells, $\rho_L = 3.56 \times 10^{-7}$–$1.967 \times 10^{-6}$ is the expression level of PD-L1 on T cells, $K_{TQ} = 1.365 \times 10^{-18}$ g/cm$^3$ is the inhibition of function of T cells by PD-L1-PD-1, and $\epsilon_C$ is the expression of PD-L1 on tumor cells versus T cells (Nikolopoulou et al (2018); Cheng et al (2013); Agata et al (1996)).

Due to parameter non-identifiability of the fraction of parameters in the denominator of (3), we define

$$\rho := \frac{\sigma \rho_P \rho_L}{K_{TQ}}, \tag{4}$$

which now represents the inhibition of T cells by formation of the PD-L1-PD-1 complex and is a scaling of the parameter $K_{TQ}$. Using the range of $\rho_P$ and $\rho_L$, we calculated $\rho$ to be $8.32 \times 10^4$–$1.22 \times 10^6$ (cm$^3$/g)$^2$. Since the units of $\rho$ are in terms of (cm$^3$/g)$^2$, similarly to the conversion of parameter $a_T$, we used the mass of a T cell (Grover et al (2011)) along with the minimum and maximum tumor volumes measured in the experimental data (Appendix B) to calculate $\rho$ in terms of cell$^{-2}$ as given in Table 1.

#### Death Rate ($d_T$)

Ribeiro et al (2002) studied the dynamics of T cells in healthy versus HIV-1 infected individuals and estimated the death rate of activated T cells. Data in (Ribeiro et al, 2002, Tab. 1) expresses a minimum T cell death rate of 0.010 day$^{-1}$ for healthy individuals and a maximum of 0.303 day$^{-1}$, which was exhibited by the untreated immune-compromised subgroup. Therefore, we anticipate the T cell death rate to be within the range of 0.010–0.303 day$^{-1}$.

#### 3.3 MDSC Parameter Values

##### Recruitment Rate of MDSCs by CCL2/CCL7 ($s_M$)

In the migration assays performed to measure MDSC recruitment to CCL2 (Takacs et al (2022)), the concentrations of chemokine (1–1000 ng/mL) used in the context of 0.15 mL of solution is equivalent to using 150 pg to 150,000 pg of CCL2. Since glioma cell expression of CCL2 ranges from 13 pg (per 10,000 glioma cells) to 111 pg (per 100,000 glioma cells), the migration assay which was most similar to glioma expression levels had 150 pg of CCL2, i.e., the assay with a concentration of 1 ng/mL. Therefore, we used the results from the 1 ng/mL assays for our calculation. Using dimensional analysis, 115,385–135,135 glioma cells would be needed to express 150 pg of CCL2.

Following nine migration assays, the average number of MDSCs that migrated to CCL2 was 229 MDSCs, while the average number of MDSCs that migrated in the control experiment (0 ng of CCL2/mL) was 91 MDSCs within 2 hours. Assuming that this implies that 138 MDSCs migrated within 2 hours specifically due to the presence of CCL2, this corresponds to a rate of 1,656 MDSCs per day. Using the range of glioma cells needed to express 150 pg of CCL2, we approximate the recruitment rate of MDSCs due to glioma expression of CCL2 to be 0.0123–0.0144 MDSCs per day per glioma cell. This calculation offers a lower bound for $s_M$ since it does not include the effect of CCL7 on MDSC migration. Note that the data for glioma cell expression of CCL2 is in terms of total concentration, thus these calculations could be improved by obtaining the extracellular concentration of CCL2 by glioma cells.

##### MDSC Expansion Coefficient ($\alpha$)

Shariatpanahi et al (2018) stated the following possibilities for the MDSC expansion coefficient, all of which were from mouse data: $0.7 \times 10^7$ cell day$^{-1}$ in EL4-luc2 lymphoma tumors (Abedi-Valugerdi et al (2016)), $1.2 \times 10^8$ cell day$^{-1}$ in 4T1 breast tumors (Cao et al (2016)), and $1.6 \times 10^7$ cell day$^{-1}$ in 3LL lung tumors (Srivastava et al (2012)). We set the range of $\alpha$ to be $1.6 \times 10^7$–$1.2 \times 10^8$ cell day$^{-1}$ since the dynamics of GBM are expected to be more similar to solid tumors rather than liquid tumors like lymphoma.

Note that these values refer to MDSC expansion in the spleen instead of the tumor site. Therefore, since we do not expect that all MDSCs in the spleen will subsequently travel to the tumor microenvironment, the relevant lower bound for $\alpha$ may be smaller.

### 4 Parameter Sets with Smallest Error

**Table S1**: Parameter sets with the smallest error (in terms of the total relative error $E_{\text{total}}$) from the ABC rejection method used to in conjunction with the minimum, average, and maximum data points.

| Parameter | Minimum | Average | Maximum |
|---|---|---|---|
| **Glioma Cells** | | | |
| $\lambda_C$ | 0.0437 | 0.431 | 0.274 |
| $C_{\max}$ | $4.18\times10^{7}$ | $3.04\times10^{6}$ | $4.20\times10^{6}$ |
| $\eta$ | $9.67\times10^{-7}$ | $2.57\times10^{-8}$ | $4.63\times10^{-10}$ |
| **T Cells** | | | |
| $a_T$ | $1.73\times10^{6}$ | $3.26\times10^{6}$ | $1.45\times10^{6}$ |
| $s_T$ | $1.16\times10^{6}$ | $8.56\times10^{6}$ | $1.91\times10^{6}$ |
| $\rho$ | 0.492 | 0.107 | $7.36\times10^{-3}$ |
| $\epsilon_C$ | 88.4 | 16.2 | 21.1 |
| $r$ | $4.28\times10^{-5}$ | $6.92\times10^{-6}$ | $9.76\times10^{-6}$ |
| $d_T$ | 0.225 | 0.0221 | 0.149 |
| **Myeloid-Derived Suppressor Cells** | | | |
| $s_M$ | $2.47\times10^{-3}$ | 0.0372 | 0.0619 |
| $\alpha$ | $3.59\times10^{8}$ | $3.60\times10^{8}$ | $4.22\times10^{8}$ |
| $q$ | $6.89\times10^{10}$ | $3.51\times10^{10}$ | $9.43\times10^{10}$ |
| $d_M$ | 0.210 | 0.251 | 0.360 |
| **Error ($E_{\text{total}}$)** | 1.78 | 0.785 | 0.937 |

### 5 Additional Bifurcation Diagrams

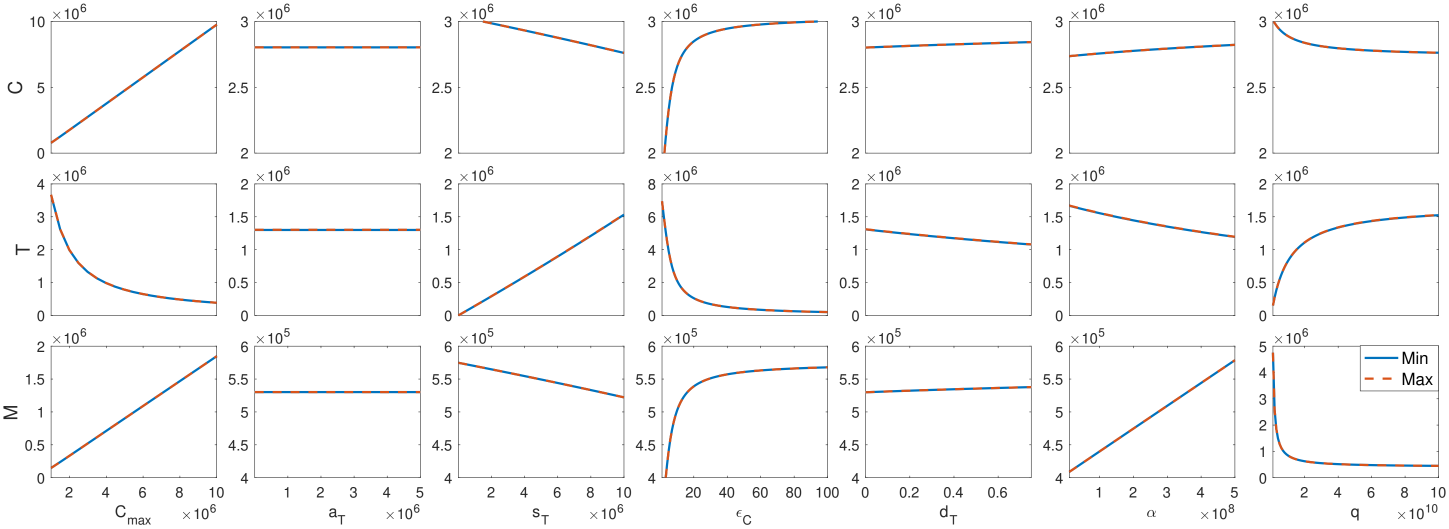

**Fig. S1**: The minimum/maximum tumor ($C$), T cell ($T$), and MDSC ($M$) cell counts are shown in solid blue/dashed red, respectively.

### 6 Additional eFAST Details and Figures

#### 6.1 Number of Samples ($N_s$) and Resamplings ($N_r$)

Since we have 13 parameters, the minimum allowable value of $\omega_{\max}$ is 13, which results in a minimum sample size of 105. However, we can improve accuracy by choosing a larger $\omega_{\max}$ in order to increase the distance between the frequencies of the complementary set $\{\omega_{(-i)}\}$. This allows us to better obtain the spectrum of the system's Fourier series due to each frequency, i.e., we can better obtain the variance of the system due to each parameter. Saltelli et al (1999), Fig. 9, compared the accuracy of several samples sizes for eFAST to analytical results for total sensitivity indices ($S_{Ti}$) and found that a sample size of roughly 2048 (including resamplings) was more than sufficiently accurate. Therefore, we generated 2049 parameter sets for each of our resamplings. Since the sample size for each resampling is determined by

$$N_s = 2M\omega_{\max} + 1, \tag{5}$$

where M is the interference factor (usually 4), the maximum frequency, $\omega_{\max}$, used was 256. According to the algorithm for the selection of complementary frequencies proposed by Saltelli et al (1999), the maximum allowable frequency for the complementary set was

$$\max\{\omega_{(-i)}\} = (1/M)(\omega_{\max}/2) = 32. \tag{6}$$

The complementary set was then chosen within the range of integers between 1 and 32 in order to maximize the step size between each frequency (in this case, the step size was 2). From experience, Saltelli et al (1999) also advised that the number of resamplings, $N_r$, be chosen in order to keep $\omega_{\max}/N_r$ within the range of 16 and 64. So, we chose to have 5 resamplings ($\omega_{\max}/N_r = 51.2$) where the phase shifts, $\varphi_i$, for each resampling was randomly chosen from $[0, 2\pi]$. Thus, this sampling method generated 2049 parameter sets for each of our 5 resamplings, resulting in a total sample size of 10,245 parameter sets.

#### 6.2 Additional Figures

In this section, we first include eFAST results at days 5, 10, and 20 in Fig. S2.

We also present a comparison between the right-skewed and uniform sampling scheme in Fig. 5 and S3, respectively, to show that the right-skewed sampling scheme narrows the search for therapeutic targets for our model.

When we compare Fig. 5 and S3, the uniform sampling method in Fig. S3 shows more interaction between parameters for all three cell populations but especially for T cells. This would suggest that many parameters need to be targeted therapeutically to control the populations. Fig. 5 thus helps us to narrow our potential targets by showcasing less interactions between parameters and highlighting only the essential ones.

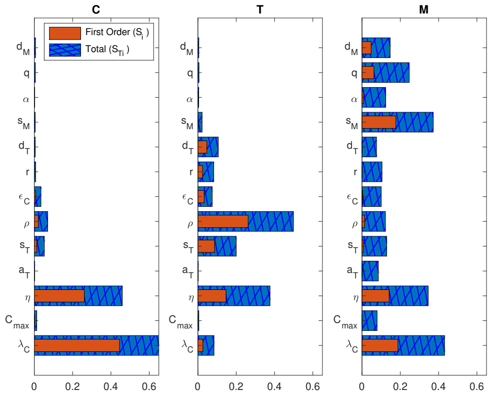

(a) day 5

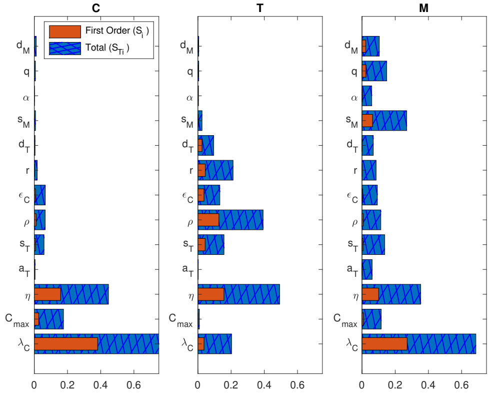

(b) day 10

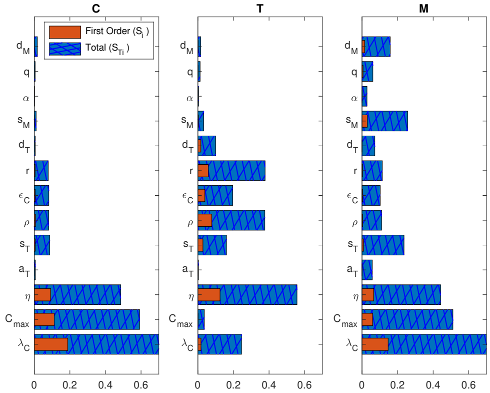

(c) day 20

**Fig. S2**: Global sensitivity analysis: The eFAST method was used to calculate the main effect $S_i$ and total effect $S_{Ti}$ of each parameter on tumor cells ($C$), T cells ($T$), and MDSCs ($M$) at different days after tumor implantation. Parameters $\lambda_C$, $C_{\max}$, $\eta$, $\rho$, $\epsilon_C$, and $r$ were sampled using (50), while the remaining were sampled uniformly using (51).

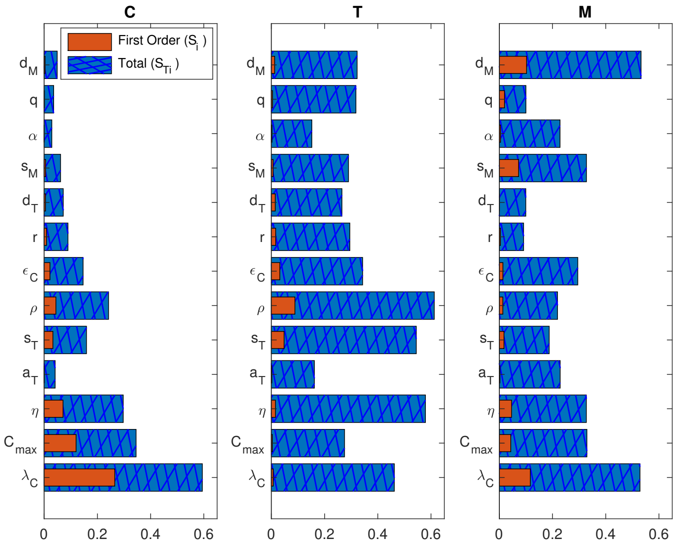

**Fig. S3**: Global sensitivity analysis: The eFAST method was used to calculate the main effect $S_i$ and total effect $S_{Ti}$ of each parameter on tumor cells ($C$), T cells ($T$), and MDSCs ($M$) at $t = 40$ days. Each parameter was sampled uniformly using (51).
